# Causal Inference on Neuroimaging Data with Mendelian Randomisation

**DOI:** 10.1101/2022.03.17.484732

**Authors:** Bernd Taschler, Stephen M. Smith, Thomas E. Nichols

## Abstract

While population-scale neuroimaging studies offer the promise of discovery and characterisation of subtle risk factors, massive sample sizes increase the power for both meaningful associations and those attributable to confounds. This motivates the need for causal modelling of observational data that goes beyond statements of association and towards deeper understanding of complex relationships between individual traits and phenotypes, clinical biomarkers, genetic variation, and brain-related measures of health. Mendelian randomisation (MR) presents a way to obtain causal inference on the basis of genetic data and explicit assumptions about the relationship between genetic variables, exposure and outcome. In this work, we provide an introduction to and overview of causal inference methods based on Mendelian randomisation, with examples involving imaging-derived phenotypes from UK Biobank to make these methods accessible to neuroimaging researchers. We motivate the use of MR techniques, lay out the underlying assumptions, introduce common MR methods and focus on several scenarios in which modelling assumptions are potentially violated, resulting in biased effect estimates. Importantly, we give a detailed account of necessary steps to increase the reliability of MR results with rigorous sensitivity analyses.

## 1 Introduction

There is an ever-present need to establish a causal interpretation for scientific data. For example, determining whether a medical intervention, such as a drug treatment, is the origin of an observed difference or change in health measures; confirming whether an environmental exposure or behavioural factor increases disease risk; or establishing whether individual traits and phenotypes contribute to adverse health outcomes, questions of causality are at the heart of scientific understanding.

Randomised controlled trials (RCTs) are considered the gold standard for inferring causal relationships, as random assignment of treatment minimises the risk of confounds causing an outcome of interest. However, RCTs are in many cases impractical or impossible, and we must depend on analytic methods that impose assumptions to bridge the gap between observational exposure–outcome associations and causal conclusions. Many of these methods rely on regression analyses and graph diagrams to infer causal relationships. Bayesian networks and related advances in graph theory, structural equation modelling and counterfactuals are some of the most prominent approaches to causal inference (Pearl, 2009; Pearl et al., 2016; Hernán and Robins, 2020).

Causal conclusions cannot safely be drawn from observational data without strong additional assumptions. An observed association between two variables of interest can be due to a true causal mechanism (in either direction), but also may arise because of an unmeasured common cause or due to sampling bias.

Mendelian randomisation (MR) presents a way to obtain causal inference on the basis of genetic data and explicit assumptions about the relationship between genetic variables, exposure and outcome. Importantly, unlike standard regression models, MR aims to be unaffected by confounding^1^ of the exposure–outcome relationship, thus excluding one of the main sources of non-causal associations in other methods.

Neuroimaging datasets on the scale of 1,000’s or 10,000’s of participants allow for population-level inquiry of disease aetiology, risk factors and biological mechanisms as they relate to the structure and function of the brain. In an aging population, the need for information on causal factors that impact brain health is apparent, especially as brain health may be a more sensitive outcome than other phenotypes. With obvious ethical and practical limitations on RCTs and interventional experiments, large-*N* population imaging is our best promise so far to identify associations between modifiable risk factors and brain phenotypes. However, massive sample sizes mean analyses are sensitive to both meaningful associations and those attributable to confounds. Conditioning on (“regressing-out”) potential confounders is often insufficient to remove non-causal associations and can even introduce additional bias (for example, via colliders, see Section 2.4. Hence, there is an urgent need for methods like Mendelian randomisation to go beyond descriptive accounts of associations and establish true causal relationships that are not the result of hidden confounding. At the same time, rigorous assessment and careful interpretation of findings from MR studies – as well as any other causal claims – are essential in order to draw plausible and valid scientific conclusions from these analyses.

For the last 20 years, Mendelian randomisation has mostly been applied to epidemiological settings. With the increasing availability of genetic data from genome-wide association studies (GWAS), the first preprints and papers using MR on neuroimaging data are now being published. Several recent studies have investigated causal links between imaging-derived phenotypes (IDPs) and various disease pathologies such as Alzheimer’s disease (Knutson et al., 2020; Garfield et al., 2020; Korologou-Linden et al., 2020, 2021; Fani et al., 2021; Wu et al., 2021), heart disease (Tian et al., 2021), depression (Shen et al., 2020), schizophrenia (Stauffer et al., 2021), other psychiatric disorders (Guo et al., 2021; Song et al., 2021) and lifestyle factors such as smoking and alcohol consumption (Logtenberg et al., 2021).

The purpose of this work is to provide an introduction to causal inference using methods based on Mendelian randomisation, with examples and background to make these methods accessible to a neuroimaging researcher. We first motivate the use of MR techniques, lay out the underlying assumptions, introduce common MR methods and give a detailed account of important steps to increase the reliability of MR results with a rigorous sensitivity analysis. Brief sections cover commonalities and differences with two other causal inference methods, mediation analysis and Bayesian networks, and how they could be used in conjunction with MR. The next section discusses several scenarios in which modelling assumptions are potentially violated resulting in biased effect estimates.

In the second part, we consider three examples focusing on the application of MR to neuroimaging data; specifically, causal relationships of systolic blood pressure, bone mineral density, and a cognitive trait with a wide range of IDPs in UK Biobank.

Going forward, we refer to traits, phenotypes and any other (risk) factors that are considered a potential cause or origin of an effect as “exposures.” Analogously, any traits, phenotypes or other factors that are potentially causally affected by an exposure are referred to as “outcomes.”

## 2 Methods

### 2.1 Mendelian randomisation

We first give a review of Mendelian randomisation before introducing other related methods that attempt to make causal inferences, namely mediation analysis and Bayesian Networks. The use of MR has grown steadily, due in part to greater availability of large-scale GWAS. In the following, we provide a high-level introduction to MR. For a detailed study we recommend recent reviews (Bowden and Holmes, 2019; Lawlor et al., 2019; Tin and Köttgen, 2021; Sanderson et al., 2022) and the comprehensive textbook by Burgess and Thompson (2015a).

Mendelian randomisation is based on the principle of using genetic variants as “instrumental variables” (see Figure 1) to investigate causal relationships in observational data (Davey Smith and Ebrahim, 2003, 2004). Instrumental variable analysis is an established methodology in the fields of econometrics, statistics and epidemiology (Lawlor et al., 2008). In addition to exposure and outcome, an instrument is a third variable that influences the outcome exclusively via its effect on the exposure. Schematically, in the causal chain *Z → X → Y* , the variable *Z* is an instrument for the *X−Y* relationship.

**Figure 1:**
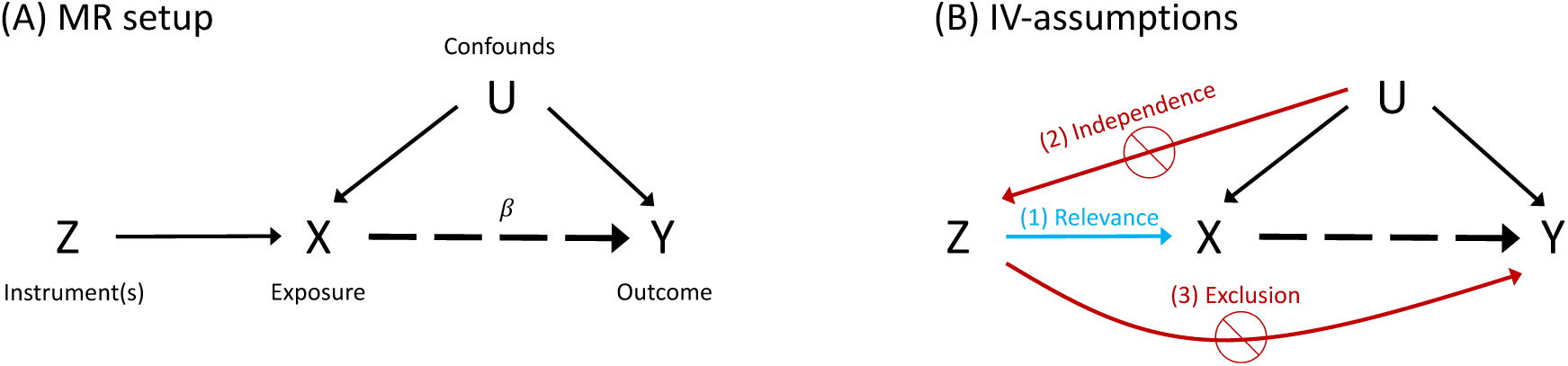
Schematic modelling of assumptions: (A) Mendelian randomisation framework, where an instrumental variable (*Z*) influences only the exposure (*X*), allowing inference on causal influence of exposure on outcome (*Y* ) even in the presence of confounds (*U* ); the causal effect of interest is indicated by the dashed arrow and *β* denotes the causal effect estimate. (B) Instrumental variable assumptions: (1) The relevance assumption, that the IV is associated with the exposure; (2) the independence assumption, that there are no unmeasured confounds of IV and outcome; and (3) the exclusion restriction, that the IV is only associated with the outcome via the exposure.

For example, consider the question of impact of alcohol consumption on liver health. There are many other factors that can influence both the level of alcohol intake and risk for liver disease, such as general health, diet, exercise and level of education. Additionally, it could be the case that liver disease affects alcohol intake. The availability of alcoholic beverages (across different countries or due to different levels of taxation), however, provides an instrumental variable that influences the chances of an individual consuming alcohol but has no direct effect on liver health. Therefore, observational data on alcohol consumption as predicted by availability can be associated with measures of liver health to obtain a less confounded estimate of the effect of alcohol on the liver.^2^ As in most scenarios, there are caveats and limitations to consider in this example. For instance, socio-economic variables will play a non-trivial role for health outcomes, levels of alcohol consumption and impact of taxation. Careful consideration of potential biases and validity of assumptions is therefore necessary.

In the remainder of this work, we will use the following example concerning the effect of blood pressure on cardiovascular health: A correlation between higher blood pressure and coronary heart disease does not prove a causal effect of one on the other, since common causes (BMI for instance) may influence both blood pressure levels and risk of heart disease. The inclusion of a new, instrumental variable, that is causally linked to blood pressure (for example, a specific genetic variant), but has no direct influence on the outcome, allows one to separate true causal effects of blood pressure on heart disease from spurious correlations due to BMI and other confounds.

In MR, the validity of the instrumental variables rests on the laws of Mendelian inheritance, in particular the principle of random assortment of parent alleles during meiosis. The fact that the composition of the genetic code is fixed at conception precludes any environmental influences or effects of lifestyle factors. This means that genetic variants are (mostly) unaffected by issues of confounding and reverse causation. In other words, comparing groups of individuals with a different genetic makeup at specific locations of the genome (e.g., single-nucleotide polymorphisms or SNPs) provides a chance to detect a causal effect between genetically-determined levels of the exposure and an outcome of interest, without many of the limitations in other observational studies that are due to confounding effects^3^. MR examines the observed association between outcome and genotype-predicted exposure. Because of the inmutability of an individual’s genotype, any robustly identified effect can be directly attributed to the exposure.

It should be noted that, although largely valid, there exist some caveats to the assumption of a fully random distribution of genetic variants among the population (see also Section 2.4). For a thorough discussion of the “MR-as-nature’s-randomised-controlled-trial” analogy and its limitations, see Swanson et al. (2017).

#### 2.1.1 Instrumental variables

For any instrumental variable (IV) analysis, there are three main assumptions that a candidate IV must satisfy to be a valid IV (Haycock et al., 2016; Burgess and Thompson, 2015a): 1) The IV is associated with the exposure (relevance assumption); 2) There are no unmeasured confounders of the association between IV and outcome (independence assumption); 3) The IV is only associated with the outcome via the exposure (exclusion restriction). A schematic summary of the standard MR setup and the three IV assumptions is depicted in the causal diagrams in Figure 1.

As an illustrative example, consider the SNP rs35479618, which has been found to be strongly associated with systolic blood pressure (Liu et al., 2016). In a uni-variable analysis, this single SNP is the instrument (*Z*), systolic blood pressure (BP) is the exposure (*X*) and coronary artery disease (CAD) is the outcome (*Y* ). Absent any direct associations of the SNP with confounds and the outcome, the simplest MR estimate for the causal effect of BP on CAD is given by the ratio of the SNP–outcome association to the SNP–exposure association. Concretely, using GWAS data on BP (GWAS ID: ukb-b-20175 (Mitchell et al., 2019)) and CAD (GWAS ID: ebi-a-GCST005195 (Van Der Harst and Verweij, 2018)), both accessed via the MRC IEU OpenGWAS data infrastructure (Elsworth et al., 2020), the SNP–BP association is 0.0617 (i.e., 0.0617 SD change in BP associated with change in SNP dosage) and the SNP–CAD association is 0.0652 (odds ratio change associated with change in SNP dosage). The causal effect of BP on coronary artery disease is therefore *β*=0.0652*/*0.0617=1.06. Since CAD is a binary variable, the effect estimate is given as a log odds ratio of CAD occurring for a one-standard-deviation increase in BP. In binary case–control scenarios, log-linear or logistic regression models are often preferred, where the effect estimate then corresponds to the log relative risk or log-odds ratio, respectively (Burgess et al., 2017b). However, due to small effects of SNPs, linear models generally approximate logistic models well and are therefore widely used in MR analyses.

By inference methods we describe below, a p-value can be computed, here p=0.017; although nominally significant, the 95% confidence interval for the effect estimate (95% CI [0.19, 1.94]) is very large. Multi-variable analyses that simultaneously use many SNPs as instrumental variables generally have higher power to detect an effect and allow for the application of more advanced MR methods as well as sensitivity analyses.

Crucially, the validity of the instrumental variable assumptions is a necessary prerequisite for the causal interpretation of Mendelian randomisation results. In practice, a potential violation of the second and third assumption can often not be ruled out and causal conclusions need to be drawn carefully. However, there are an increasing number of sensitivity analyses as well as robust MR methods available that can aid in the identification of bias, and support tentative causal claims (see Section 2.1.5).

#### 2.1.2 Individual- vs. summary-level data

MR can be performed using individual subject-level data or summary statistics from large-scale genome-wide association studies (i.e., regression coefficients and standard errors of the SNP–phenotype associations). Although, conceptually, the two approaches are equivalent, in practice, each has its own benefits and drawbacks. Individual-level MR allows one to test and adjust for suspected SNP–confounder associations and to perform subgroup analyses, but usually has lower statistical power to detect causal effects due to smaller sample sizes. Summary statistics from international GWAS consortia on the other hand are readily available and often based on very large sample sizes, and are commonly used in so-called two-sample MR, where the SNP–exposure and SNP–outcome associations are estimated on two separate datasets (Burgess et al., 2015). Individual-level data is often used in one-sample MR, which is more prone to overfitting due to weak instrument bias (see Section 2.4). One-sample settings have the potential benefit that MR results can be linked to other analyses involving the same individuals, whereas two-sample analyses would be problematic if the two data sets differ substantially in their population characteristics (ethnicity, sex, age, socio-economic status, etc.) (Burgess et al., 2020a).

Because of potential bias due to sample overlap and weak instruments, two-sample MR is commonly preferred in practice. However, the two datasets in two-sample MR must represent the same population, and summary effect estimates need to be harmonised across the two datasets.

Some specialised MR approaches are only available for individual-level data, such as factorial MR to assess interactions (Rees et al., 2020) and non-linear MR (Staley and Burgess, 2017; Silverwood et al., 2014). Recent advances in methodology continue to expand the availability of MR variants to summary-level data, for example, methods for identifying violations of the exclusion restriction, known as horizontal pleiotropy, via gene-by-environment interactions (Spiller et al., 2019).

For the remainder of this paper, we will focus on summary-level MR methods since these are i) more common, ii) easier to carry out, and iii) as far as neuroimaging phenotypes are concerned, summary statistics from population studies such as UK Biobank are essentially the only available data with large enough sample sizes (ideally, *N ≫* 10^4^).

#### 2.1.3 SNP selection and pre-processing

Genetic variants are usually selected based on a significance threshold of the SNP–exposure association from GWAS results (typically *p <* 5*×*10*^−^*^8^, but might have to be adjusted based on sample size). For a single SNP, this p-value is an indirect measure of the effect size but crucially also depends on overall sample size and frequency of occurrence (the minor allele frequency or MAF) of the genetic variant in the sampled data (Swerdlow et al., 2016). If prior knowledge is available on which gene or gene region is implicated in the regulation of the exposure of interest, then the selection of genetic variants can be restricted to that region of the genome only. This approach has been used successfully, for example, to determine the causal effect of LDL cholesterol on coronary heart disease while ruling out the HDL variant (Schmidt et al., 2020; van der Graaf et al., 2020). Otherwise, a polygenic analysis involving genetic variants from potentially multiple genetic regions is performed. Most robust MR methods assume independence between SNPs, and thus it is important to have SNPs sufficiently separated in genetic distance (Burgess et al., 2020a). Additionally, the inclusion of polygenic variants that explain independent parts of the exposure–variance (i.e., with different biological pathways from the genetic variants to the exposure) improves the statistical power to detect a causal effect.

Prominent metrics for instrument selection include the proportion of variance explained (*R*^2^) and the F-statistic^4^ of the exposure-on-SNP regression model (Burgess and Thompson, 2011; Swerdlow et al., 2016). A threshold of *F >* 10 is conventionally considered as an indicator for sufficiently strong instruments.

MR analyses that involve more than a single genetic variant require the clumping of SNPs as a preprocessing step. This ensures that the SNPs used as instrumental variables for the exposure are independent. SNPs with allele frequencies that vary together to a degree outside of what would be expected from a random, independent association are considered to be in linkage disequilibrium (LD). A reference database such as the 1000 Genomes reference panel (Altshuler et al., 2010) can be used to calculate LD *R*^2^ values for a set of selected SNPs. Above a certain cutoff (typically *R*^2^ *>* 0.001), only the SNP with the lowest p-value for the SNP–exposure association is retained, thus “clumping” together (though in actuality, discarding) covarying SNPs.

In two-sample MR, care must be taken to ensure that the GWAS-reported effect of a selected SNP on the exposure (in one dataset) and the reported effect of the same SNP on the outcome (in another dataset) correspond to the same allele. Updates to the human genome reference sequence as well as changes in the way GWAS data are reported mean that SNP annotation often differs between genotyping platforms, datasets and repositories, making mismatches a common problem. MR software such as the `TwoSampleMR` (Hemani et al., 2018b) and `MendelianRandomization` (Yavorska and Burgess, 2017) R packages include harmonisation procedures that can infer the correct allele alignment as automated preprocessing steps.

#### 2.1.4 Standard MR methods

The causal effect estimate for the elementary case based on summary-level data and a single SNP is simply given by the ratio of the SNP–outcome association to the SNP–exposure association^5^. From the regression model for the SNP–exposure association *γ_E_* on the SNP–outcome association *γ_O_*, we have 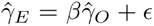, where *ɛ* denotes the error term; the effect estimate is then obtained as 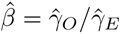^6^. When multiple SNPs are available, the ratio estimates for each SNP can be combined in a meta-analytic fashion to estimate an overall causal effect (Burgess et al., 2013; Bowden et al., 2016a). This constitutes the standard inverse-variance weighted (IVW) MR method.

Several meta-analytic approaches are possible, including fixed-effects, additive random-effects and multiplicative random-effects models. Their suitability is determined by the presence of heterogeneity and pleiotropy. In the presence of balanced pleiotropy, that is when positive and negative pleiotropic effects on the outcome on average cancel out, fixed-effects and additive random-effects meta-analyses are both unbiased. A difference in their respective effect estimates indicates directional (unbalanced) pleiotropy, in which case a fixed-effects model is preferred. If strong heterogeneity (large variance in individual SNP estimates) is detected, an additive random-effects model will give greater weight to weaker (and more biased) single-SNP estimates and a multiplicative random-effects model is recommended (Burgess et al., 2020a). In a multiplicative random-effects model, heterogeneity in the single-SNP estimates does not influence the point estimate 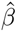. However, the variance of 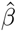 is allowed to increase with heterogeneity. Multiplicative random-effects models are also known to be more robust to small sample bias. In practice, these three IVW MR variants can be used as part of a sensitivity analysis. Large differences in their causal effect estimates are a sign of (directional) pleiotropy or problematic heterogeneity. An in-depth account of meta-analytic approaches for MR can be found in Bowden et al. (2017).

Although it is the most efficient method in terms of statistical power, standard IVW MR is not robust to outliers and requires all selected SNPs to be valid instrumental variables. In IVW MR, the intercept is fixed at zero (see Figure 3B). This follows directly from the third IV-assumption that all selected SNPs are acting on the outcome only via the exposure. Thus a null association with the exposure entails a zero effect on the outcome. MR-Egger regression (Bowden et al., 2015; Burgess and Thompson, 2017) removes this constraint, allowing the intercept to vary freely. Consequentially, a non-zero estimate for the MR-Egger intercept indicates the presence of invalid instruments due to pleiotropy, and hence allows this violation of the MR assumptions to be flagged (Burgess et al., 2020a). Although MR-Egger permits all instruments to be affected by pleiotropy, the so-called InSIDE (Instrument Strength Independent of Direct Effect) assumption requires that any pleiotropic effects are independent of the instrument–exposure associations. One of the main drawbacks of the MR-Egger approach are its high sensitivity to outliers and its reduced efficiency (lower power) compared to IVW MR.

Many robust MR methods rely on relaxed assumptions for a subset of instrumental variables in a polygenic analysis framework. These methods can usually deal with some fraction of invalid instruments and still provide valid causal inferences. While problems such as linkage disequilibrium or systematic confounding due to selection bias can invalidate the analysis, instrument invalidity most often arises from horizontal pleiotropy.

This violation of the exclusion restriction, horizontal pleiotropy, arises when the genetic variant influences the outcome via additional pathways that do not include the exposure (see Figure 2D) (Burgess et al., 2020a).

**Figure 2:**
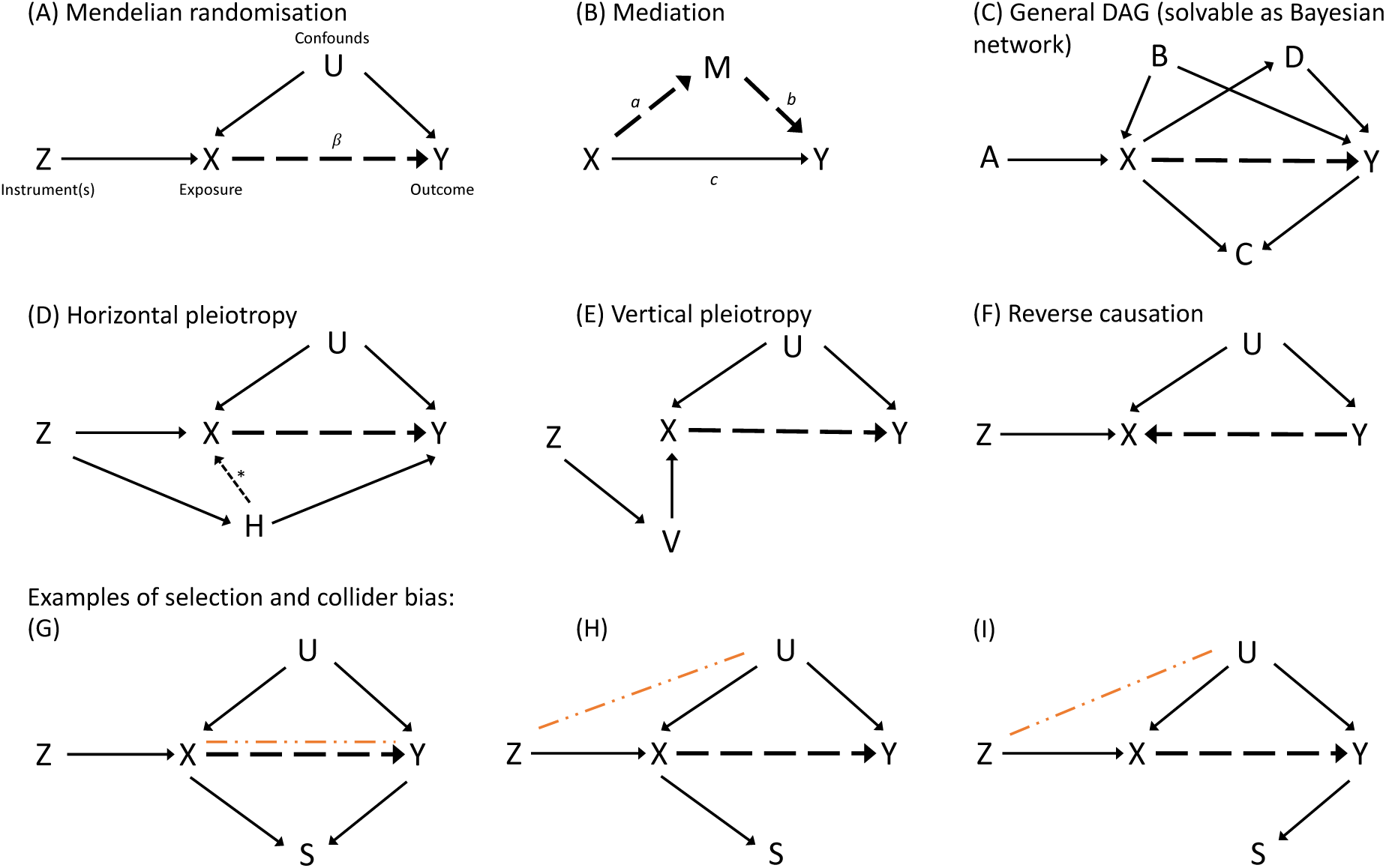
Directed acyclic graphs depicting different causal modelling approaches (A-C) and various scenarios of potential bias (D-I): (A) Mendelian randomisation. (B) Mediation. (C) General DAGs as Bayesian networks: Shown is an example DAG that includes exogenous causes *A*, a common cause *B*, a collider *C*, and a mediator *D* of the *X Y* relation. Note that this shows only one out of a large number of possible configurations for the same set of variables. (D) Violation of the exclusion restriction assumption in MR due to horizontal pleiotropy via variable *H*. The case where *H* also affects the exposure (indicated by the dashed arrow with a star next to it) is called correlated horizontal pleiotropy; otherwise it is called uncorrelated. (E) Vertical pleiotropy via variable *V* . In principle, vertical pleiotropy is not a problem for MR. However, in practice, vertical pleiotropic effects cannot easily be separated from horizontal pleiotropic effects. (F) Model mis-specification due to reverse causation. (G-I) Three cases of selection bias due to conditioning on variable *S*. The orange dot-dashed line indicates an induced association when conditioning on *S*. Panel G is an example of collider bias for the *X Y* association. Panels H and I are examples of collider bias that introduces a spurious *Z U* association, which in turn violates the IV assumptions. (Note that the presence of *S* alone is not a problem; bias only arises when conditioning on *S*.) Dashed black arrows indicate the relationship of interest.

**Figure 3:**
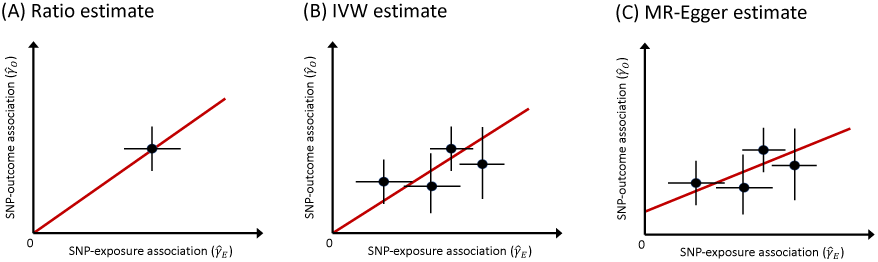
Schematic depiction of standard Mendelian randomisation models: (A) Single-SNP ratio estimate. (B) Multiple-SNP inverse variance weighted MR estimate. (C) MR-Egger estimate with non-zero intercept. The causal effect estimate is the slope of the red line, i.e., the expected increase in the SNP–outcome association for each unit increase in the SNP–exposure association. Errorbars indicate standard errors for each SNP association. In (A) and (B) the intercept is assumed fixed at zero.

Robust methods generally allow a certain number of SNPs to be affected by pleiotropic effects under the condition that the instrumental variable assumptions hold for the rest of the selected instruments. In the following we briefly cover the main characteristics of the most commonly used MR approaches.

Among robust methods, median-based (Bowden et al., 2016a) and mode-based (Hartwig et al., 2017) methods take a consensus-based approach by assuming that a majority (median-based) or a plurality (mode-based) of instruments are valid. By using, in effect, only a subset of instruments, these methods are more robust to outliers in the single-SNP ratio estimates of the causal effect at the cost of reduced statistical power.

Another approach, MR-PRESSO (Verbanck et al., 2018) allows for up to half of selected instruments to be affected by horizontal pleiotropy. It seeks to find and remove outliers based on a heterogeneity test of the single-SNP effect estimates. After removal of genetic variants with substantially different effect estimates, a standard IVW MR analysis is performed. The reason for demanding all instruments to give similar effect estimates is that this should reflect the same causal effect of the exposure on the outcome, regardless of the choice of IV and the strength of the IV’s causality onto exposure and outcome.

In scenarios where several, closely related exposures are candidate causes for the same outcome, genetic variants that are selected as instrumental variables may be associated with some or all related exposures. This means that it may not be possible to find a set of instruments that is specific to one exposure without exhibiting pleiotropic effects via related exposures. Multivariable MR (MVMR) approaches (Burgess and Thompson, 2015b; Sanderson et al., 2019) try to address this by including multiple exposures in the analysis. The standard IV assumptions must still hold, with the set of exposures replacing the single exposure in univariable MR. Conceptually, multivariable MR is an extension of the IVW method and estimates a causal effect while controlling for a number of other, measured biological pathways.

Overall, MR methods development is an active area of research and many novel or modified analysis approaches have been proposed in recent months and years. A full account is beyond the scope of this paper, however, many newer methods fall into one of two broad categories: i) regression models (e.g., MR-Lasso (Slob and Burgess, 2020), radial regression (Bowden et al., 2018), and ii) likelihood-based models (e.g., MR-Mix (Qi and Chatterjee, 2019), MR-RAPS (Zhao et al., 2020), BayesMR (Bucur et al., 2020), contamination mixture (Burgess et al., 2020b), CAUSE (Morrison et al., 2020), GRAPPLE (Wang et al., 2020)).

For an extensive overview of standard and robust MR methods and best practice approaches, we highly recommend recently published guidelines (Burgess et al., 2020a; Davey Smith et al., 2019; Sanderson et al., 2022), method comparisons (Slob and Burgess, 2020) and the STROBE-MR guidelines on reporting MR results (Skrivankova et al., 2021).

#### 2.1.5 Sensitivity analysis of MR results

Recent reviews and guidelines (Burgess et al., 2020a; Davey Smith et al., 2019) strongly advocate for the inclusion of sensitivity analyses as a core part of any Mendelian randomisation investigation.

The main approach to assess the robustness of findings from MR analyses is to obtain several estimates from different MR variants, including standard mean and median based methods, MR-Egger regression, methods sensitive to outliers such as MR-PRESSO, multivariable MR, and any other approach that may be suitable for the particular data at hand. An additional, straightforward assessment can be done by varying the selection of instruments via a more (or less) stringent p-value threshold for the SNP–exposure association. Consistency of results from different sets of genetic instruments with fewer but stronger (or more but weaker) instruments will generally indicate a robust causal effect.

A similar approach is to look for heterogeneity in effect estimates as a result of removing a single instrument or a subset of instruments from the analysis. Leave-one-out and SNP-subset analyses can help identify variants that are predominantly driving the causal effect estimate. In cases where a bi-directional causal pathway may exist between exposure and outcome or when the direction of the causal effect is part of the research question, Steiger filtering (Hemani et al., 2017) can be used to remove potentially invalid genetic variants under the assumption that the SNP–exposure association is expected to be stronger than the SNP–outcome association. However, Steiger filtering is sensitive to measurement errors and may lead to the removal of valid instruments, especially in two-sample MR analyses.

Further recommended is the inclusion of heterogeneity measures such as Cochran’s *Q* statistic or the *I*^2^ statistic^7^ in polygenic MR analyses (Bowden et al., 2016b). Substantial heterogeneity in SNP-specific causal effect estimates and clear outliers can indicate the presence of horizontal pleiotropic effects. On the other hand, largely homogeneous effect estimates provide the basis for more reliable causal conclusions (Burgess et al., 2020a).

Various graphical tools can be used to gain qualitative information about potential outliers and unexpectedly skewed or otherwise biased effect estimates. These include scatter plots of the SNP associations with exposure and outcome, funnel plots of single-SNP effect estimates and forest plots of leave-one-out analyses (see, e.g., Figures 5, 13).

When prior knowledge about causal relationships involving the exposure and outcome variables is available, a positive outcome or negative outcome analysis can help establish the validity of chosen instrumental variables (Sanderson et al., 2021a). For example, using a positive control outcome and given a large enough sample size, SNPs that do not yield an effect similar to what has already been established may be too weak or invalid for the exposure in question and are unlikely to produce correct effect estimates when used with the outcome of interest.

A summary of suggested steps and procedures to be considered when performing an MR analysis is provided in Table 1.

**Table 1:** Summary of essential steps in a comprehensi^1^v^1^e MR and sensitivity analysis. Items marked by **\*** are essential or strongly recommended.

| (0) DATA COLLECTION |  |  |
| --- | --- | --- |
| What | Why | How |
| Individual-level data * | One-sample MR | Genome-wide (or at least instrument-specific) genotype data |
| OR |  |  |
| Summary statistics (effect sizes, standard errors) * | One-sample or two-sample MR | GWAS databases (e.g., IEU Open GWAS project, EBI) |
| SNP annotations | Biological interpretation | Reference databases |
| (1) DATA PREPARATION |  |  |
| What | Why | How |
| Clumping and harmonisation of genetic variants * | Instrument validity | Standard MR tools |
| Proportion of variance explained in exposure ($R^2$) | Assessment of instrument strength | Standard statistical tests |
| Mean F-statistic of regressors (recommended $\bar{F} > 10$) | Assessment of instrument strength | Statistics; for MVMR see Sanderson et al. (2021b) |
| Varying IV–exposure association threshold | Robust effect estimate | Standard MR tools |
| (2) MR ANALYSIS |  |  |
| What | Why | How |
| IVW MR * | Standard, most efficient effect estimate | Standard MR tools |
| MR-Egger * | Robust effect estimate | Standard MR tools |
| Median- and/or mode-based MR * | Robust effect estimate | Standard MR tools |
| MR-PRESSO * | Heterogeneity and outlier detection, robust effect estimate | MR-PRESSO R package |
| MR-Mix | Robust effect estimate | MRMix R package |
| multivariable MR | In case of multiple related exposures | MVMR R package |
| Any additional robust / novel MR methods | Different underlying assumptions, triangulation of evidence | See, for example, methods listed in Sanderson et al. (2022) |
| Bi-directional MR * | In case of potential reverse causation | Standard MR tools |
| (3) SENSITIVITY ANALYSIS |  |  |
| What | Why | How |
| MR-Egger intercept * | Heterogeneity detection | Standard MR tools |
| Cochran’s $Q$, Higgins’ $I^2$ statistic * | Heterogeneity detection | Statistics |
| Steiger filtering | Reverse causation | Standard MR tools |
| Meta-analytic IVW MR variants | Heterogeneity detection | Standard MR tools |
| Leave-one-out or IV-subset analysis * | Heterogeneity detection | Standard MR tools |
| Single-SNP analysis | Heterogeneity detection | Standard MR tools |
| MR-RAPS | Outlier detection | mr.raps R package |
| Radial-MR | Outlier detection | RadialMR R package |
| Scatter plot of IV–outcome vs. IV–exposure associations * | Heterogeneity and outlier detection | Plotting tools |
| Funnel, forest and radial plots of individual effect estimates * | Heterogeneity and outlier detection | Plotting tools |
| Prior knowledge about biological mechanisms | Validity of results | Literature |
| Triangulation of evidence * | Reliability of results | Literature, complementary methods |

#### 2.1.6 Interpretation of results

Depending on the nature of the investigation, a distinction can be made whether the presence of a causal effect and its direction (testing the causal null hypothesis) is of primary interest or whether the goal is an estimation of the effect size. Scenarios in which the expected effects of an intervention (e.g., drug treatment or medical procedure) on the exposure are scrutinised are mostly concerned with effect size, whereas questions of disease aetiology or fundamental biological mechanisms focus predominantly on causal direction detection.

In addition to the three core IV assumptions discussed in Section 2.1.1, an additional monotonicity or homogeneity assumption needs to hold for a causal interpretation of the MR effect estimate. Monotonicity refers to the relationship between genetic instruments and exposure. It assumes that the SNPs could not increase the level of exposure in some individuals and decrease the level of exposure in others. Homogeneity is a slightly stronger assumption in that it requires that the effect of the SNPs on the exposure (or the effect of the exposure on the outcome) is the same for all individuals. If any of these two assumptions hold then the MR estimate is consistent with the average causal effect for the population under study (Burgess and Thompson, 2017; Swanson et al., 2018).

In the context of Mendelian randomisation, the causal effect of the risk factor on the outcome is commonly interpreted as the consequence of a lifetime exposure to a genetically determined level of the risk factor. The impact as well as the biological pathways through which the risk factor influences the outcome may be different in case of a direct intervention. Under valid instrumental variable assumptions, an MR estimate can be regarded as the causal effect when determining someone’s genotype at conception. For the purposes of modification and intervention in clinical settings, additionally an equivalence between gene and environment effects needs to hold.

This is further complicated when time-varying exposures are considered. In these cases the estimated effect from MR should be interpreted as the effect of changing the (genotypic) liability that causes the exposure as a function of time (Morris et al., 2021). Recommendations from methodological researchers strongly advise against a simplistic interpretation of effect estimates and emphasise the perspective that MR should be used to test the causal null hypothesis rather than to estimate effect magnitudes (Vanderweele et al., 2014; Burgess et al., 2020a, 2021).

#### 2.1.7 Limitations

Apart from limitations inherent in the modelling assumptions underpinning Mendelian randomisation, there are issues that can arise from the data itself, e.g., from the way data are collected and processed or in terms of sample size and composition.

Focusing on UK Biobank, several recent papers have reported findings that demonstrate non-random patterns in the data. For example, Haworth et al. (2019) have identified a geographical structure in UKB genotype data. A coincidence of health outcomes and genetic variants with birth location can introduce biased associations and potentially invalidate modelling assumptions. Other studies have shown that population structure (Lawson et al., 2020) and selection bias (Munafò et al., 2018) may play a non-negligible role in the composition of UKB samples, and that genotypic information can predict participation in some components of the UKB assessments (Tyrrell et al., 2021).

Many bias issues can be seen as different versions of selection bias (in the causal literature commonly referred to as collider bias), where two or more variables influence whether someone is selected for or takes part in a data collection study. In statistical terms, this means that selection into the sample is conditional on a common cause of the variables in question, thereby introducing a spurious association between these variables (see Figure 2G–I for a graphical representation). Selection and other biases, most notably pleiotropy in the case of MR, are discussed in Section 2.4.

### 2.2 Mediation analysis

In mediation analysis the goal is to investigate the (potentially) indirect relationship between an exposure variable and an outcome variable, where the indirect causal effect of the exposure on the outcome is mediated by a third variable. Standard mediation analysis estimates total, direct and indirect effects of the exposure on the outcome (see Figure 2B). Denote the total effect of the exposure on the outcome as *β_tot_*, the effect of exposure on mediator as *a*, and the effect of mediator on outcome as *b*. Estimates for *β_tot_*, *a* and *b* are obtained by regressing the outcome on the exposure 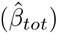, the mediator on the exposure 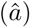, and the outcome on both exposure and mediator 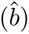, respectively. The total effect simply takes into account all potential pathways from exposure to outcome. The indirect effect accounts for the pathway from exposure to outcome via the mediator (or set of mediators) and is often the quantity of interest, for example when considering mediators as interventional targets in cases where the exposure cannot be intervened upon. The remaining effect of the exposure on the outcome that acts via pathways not including the mediator is captured by the direct effect (denoted as *c* in Figure 2B), which can be estimated by controlling for the mediator in an outcome-on-exposure regression.

Traditional methods to estimate the mediated effect (*β_med_*) are i) subtracting the direct effect from the total effect (difference method, 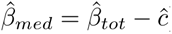), or ii) multiplying the coefficient of the exposure–mediator association with the mediator–outcome association (product of coefficients method, 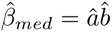). In scenarios involving only continuous outcomes and mediator variables, modelled with linear regression and fit via ordinary least squares, the two methods are asymptotically equivalent (VanderWeele, 2016).

Returning to our previous example of how blood pressure affects coronary artery disease, one can pose a related question in terms of a mediation framework. For example, one might be interested in the BP-mediated effect of body mass index (BMI) on CAD. In this scenario, BMI is the exposure, BP the mediator and CAD again the outcome. It is known that high BMI increases BP. However, BP can more easily be controlled through medication and thus some of the harmful effects of BMI on cardiovascular health can be partially controlled. A mediation analysis could try to answer the question of how much of the total effect of BMI on CAD is mediated by BP. Potential confounders of the BMI–CAD relationship, such as age, sex, smoking, physical activity, diet, etc., would also need to be included in the regression models. For a detailed mediation study of this very question, see for example Lu et al. (2015).

Standard mediation analysis relies on strong, essentially untestable assumptions regarding the absence of unmeasured confounding, exposure–mediator interactions and measurement errors. Carter et al. (2021) review two increasingly popular ways in which MR can be applied to mediation analysis in order to estimate direct and indirect effects. The first approach, multi-variable MR (MVMR) (Sanderson, 2020) treats the original exposure and the mediator (or set of mediators) as multiple exposures, using a common set of instrumental variables. In MVMR, the direct effect is estimated by controlling for the mediator, and the indirect effect can be obtained similarly as in the difference method. In the second approach, two-step MR, two separate MR analyses are performed (one for the the exposure–mediator relationship and one for the mediator–outcome relationship). The indirect effect is then estimated by forming the product of these two causal effect estimates, analogous to the product of coefficients method. For a detailed exposition of how MR techniques can be applied in a mediation framework, we refer the interested reader to the excellent review by Carter et al. (2021).

Fundamentally, Mendelian randomisation as well as mediation analysis are linear regression models. The causal interpretation of coefficients and effect estimates requires the validity of a set of strong (although often plausible) assumptions for each of the two frameworks. The difficulty lies in reasonably justifying these assumptions whenever causal claims are concerned.

### 2.3 Bayesian networks

Bayesian networks (BN) describe the conditional independence relationships of a set of variables with the help of a directed acyclic graph (DAG), which provides a graphical representation of the estimated causal structure (Figure 2C), and an accompanying joint probability distribution. Mathematically, the factorisation of the joint probability over all variables is equivalent to graphical independence properties (*d-separation*). Recent review articles (Glymour et al., 2019; Daly et al., 2011; Bielza and Larrañaga, 2014; Kyrimi et al., 2021) survey the current state and ongoing research activities regarding graphical causal modelling alongside the advances of the field over recent decades.

Various algorithms have been developed to infer the DAG that best fits the data. Broadly speaking, causal discovery algorithms can be grouped into i) constraint-based and ii) score-based methods. Constraint-based approaches start with a fully connected graph and carry out a series of marginal and conditional independence tests to iteratively remove edges that fail these tests. The PC and FCI algorithms are classic representatives of this class and are widely used in causal inference applications. Constraint-based methods estimate a Markov equivalence class, that is, a set of DAGs (often more than one) with different causal structures that satisfy the same conditional independence relations. Score-based methods, on the other hand, require a likelihood and perform a model search in graph space with the aim of optimising the score of a chosen score function, for example the Bayesian information criterion (BIC).

An important distinction has to be made between *probabilistic* and *causal* graphical models when interpreting the associations entailed in a (causal) Bayesian network (Pearl, 2009). A causal interpretation requires three key assumptions to be satisfied: i) Every variable is independent of its non-descendants conditional on its parents (causal Markov assumption); ii) There exist no other conditional independence relations other than the ones implied by the causal DAG (causal faithfulness assumption); and iii) There are no hidden common causes of two or more variables (causal sufficiency assumption). Additionally, it is assumed that there is no measurement error in the observed values of the model variables.

For a review of studies that have used Bayesian networks in healthcare in general and with neuroimaging data in particular, see for example, citeBielza2014 and Kyrimi et al. (2021). Several well established software implementations of graphical model algorithms exist, such as the bnlearn (Scutari and Denis, 2021) and pcalg (Kalisch et al., 2012) R packages.

Ancestral graphs are an extension of DAG models and are based on estimating the transitive closure of a graph (i.e. not only including an edge between a node and its direct causes but also every indirect cause or ancestor). While standard Bayesian networks estimate a direct causal effect between two variables, ancestral graph methods estimate the total causal effect on any given node.

Most standard methods require the absence of confounders, i.e., latent (non-measured) variables that are common causes of any two or more variables in the model. An exception is the FCI algorithm which produces asymptotically correct results even in the presence of confounding (Glymour et al., 2019). FCI is based on ancestral graphs and can identify spurious associations caused by latent confounding. However, due to underlying assumptions, this is restricted to scenarios that only involve jointly Gaussian distributed variables, a limitation that is unlikely to hold in the case of neuroimaging data (Grosse-Wentrup et al., 2016). Whereas a DAG consists of only directed edges, representing an association between the parent (cause) and child (effect) variable when all other variables are held constant, the representation of an equivalence class (a partially directed DAG or PDAG) can also contain undirected or bidirected edges.

Several algorithms (Kalisch and Bühlmann, 2014; Colombo et al., 2012; Zhang, 2008) have been proposed to estimate the ancestral graph structure, including FCI, RFCI, IDA and LV-IDA which are implemented in the pcalg R package.

#### 2.3.1 Bayesian networks vs. Mendelian randomisation

A recent paper (Howey et al., 2020) demonstrated that MR and BN methods can be used as complementary approaches in causal inference applications where pleiotropic effects and confounding may play a significant role. The authors showed, using both simulations and real data, that the use of directional anchors can greatly improve graphical structure estimation with standard BN algorithms, and that under certain conditions, BN approaches can give more accurate results than MR.

Unlike Mendelian randomisation, which can be used with both individual and summary-level data, Bayesian networks require individual-level data (or other sources of distributional information, e.g., covariance or conditional dependency matrices). This makes them less applicable when only summary statistics from GWAS databases are available. However, BN methods can be considered complementary to MR-based investigations, as the two approaches use very different algorithms for causal inference, rely on different sets of assumptions, produce different outputs and, when using summary-level MR methods, are based on different kinds of data.

Bayesian networks have the additional advantage that directional anchors can easily be implemented in the form of a *white-list* (and/or *black-list*) of required (excluded) edges, which can be interpreted as prior knowledge about causal (in)dependencies between specific variables. Especially for larger networks, including/excluding known edges can substantially reduce the search space of possible causal models and help identify the correct model within an equivalence class. In a typical example using SNPs as directional anchors, edges from SNPs known to be associated with a variable to the corresponding GWAS targets would be white-listed and any edges directed into SNPs black-listed. Additional domain knowledge can be used analogously to orient individual edges and exclude biologically unreasonable or impossible connections.

Bayesian network methods are limited by their strong dependence on the set of chosen measurable covariates that are included in the model. Further, scalability is an issue, preventing the inclusion of large sets of variables in most practical applications. Standard BN approaches are generally not robust against hidden confounders and require specific distributional assumptions. Importantly, the causal relationships implied by each graphical model are only strictly valid under strong (often untestable) assumptions. Mendelian randomisation (and instrumental variable analysis in general) is a dramatically different approach in that it uses the assumed (albeit very simple) causal structure given in Figure 1A to justify a projection of the problem into the space of variance explained by the instruments. Put another way, a Bayesian network will always have more variance at its disposal to estimate relationships, but at the cost that it relies on much broader assumptions than MR.

### 2.4 Common sources of bias and confounding in Mendelian randomisation

All statistical models depend on assumptions, but since the premise of Mendelian randomisation is causal inference from observational data, there is particular scrutiny on MR assumptions. Sources of bias and confounding can broadly be attributed to either i) data and study design issues (selection bias, family or dynastic effects, etc.), ii) methodological and statistical issues (weak instruments, sample overlap, small sample size, etc.) or iii) model mis-specification (e.g. pleiotropy, reverse causation).

Pleiotropy, specifically *horizontal pleiotropic effects*, i.e., pathways from genetic variants to the outcome that do not go through the exposure (Figure 2D) are a major concern in MR studies. Vertical or mediated pleiotropy (Figure 2E), that is, when the effect of the instrument on the exposure is mediated by one or more additional variables, is generally not problematic, unless the exposure in question is considered as a target for intervention and the goal of the MR analysis is to estimate the expected effect size.

Horizontal pleiotropy is often exacerbated when considering high-level phenotypes that are far removed from the level of genes and proteins (Swerdlow et al., 2016). It is generally easier to study associations earlier in the biological chain where genetic effect sizes and specificity of associations are greater. More complex phenotypes require larger sample sizes and more careful checking for possibly confounding effects. Easily obtainable indicators of horizontal pleiotropy are a non-zero intercept in the MR-Egger regression, heterogeneity tests via Cochran’s *Q* statistic, and asymmetry in the funnel plot of single-variant effect estimates. Funnel plots are commonly used in meta-analyses. They show instrument strength ( 1*/SE* denotes the inverse standard error and is a measure of precision of the estimated effect, which increases with sample size ) on the y-axis and the single instrument estimate on the x-axis, and are based on the assumption that more precise estimates are less variable, creating a triangular envelope.

Symmetry in the funnel plot indicates balanced pleiotropy, whereas asymmetry is a sign of directional (unbalanced) pleiotropy. See, for example, Figure 5B. In the presence of pleiotropic pathways, the overall causal effect may still be unbiased if, on average, positive and negative pleiotropic effects balance out (Burgess et al., 2017a). To avoid horizontal pleiotropy, MR-PRESSO and other outlier-robust methods (Slob and Burgess, 2020) should be used. Additionally, one can assess associations of exposure and outcome variables with covariates that are potentially on pleiotropic pathways.

Additionally, down-weighting or removal of outliers can provide more robust estimates (Hemani et al., 2018a). In cases where pleiotropic pathways are known, multi-variable MR can control for SNP–outcome associations via multiple exposures. Recently, MR methods have been proposed that try to accommodate certain levels of pleiotropy while still giving valid causal estimates (see, for example, Berzuini et al. (2020); Patel et al. (2021), and references therein).

In line with the IV assumptions, an instrumental variable is assumed to be marginally independent of any confounder. When conditioning on a collider of the instrument and a confounder, *selection bias* (Cole et al., 2010; Munafò et al., 2018) can lead to a spurious association between the exposure and the outcome (Figure 2G), even in the absence of any true causal relationship between exposure and outcome. Selection based on values of the exposure (Figure 2H) or values of the outcome (Figure 2I) can lead to spurious associations between the instrumental variable and the outcome via unmeasured confounders. Direction and magnitude of the bias are generally application-dependent but it has been shown that the number of false positives (type I error inflation) is more severe with larger sample sizes or very strong instruments, and that a strong dependence of the selection process on either the outcome or the exposure can have a large impact on estimated effect sizes and direction (Gkatzionis and Burgess, 2019). One way to address selection bias after data collection is via inverse-probability weighting, where each observation is weighted inversely according to its predicted probability of inclusion in the model.

*Survivor bias* (Smit et al., 2019; Schooling et al., 2021) is a particular type of selection bias where selection effects are due to mortality. This can be an issue if GWAS data is based on an older population. If it is possible to identify competing risk factors and common causes of survival and the outcome of interest, then these can be controlled in the GWAS. Alternatively, multi-variable MR or negative control outcomes can be used (Sanderson et al., 2021a). The latter would be able to detect the presence of population stratification in the GWAS of the phenotype of interest.

Bias due to *sample overlap* occurs in two-sample MR when the SNP–exposure and SNP–outcome associations are not based on completely distinct subjects. However, this issue is considered to be less problematic because any bias of the effect estimate due to sample overlap is in direction of the null (Burgess et al., 2016). For one-sample settings and overlapping samples, the estimate is asymptotically unbiased but can exhibit substantial bias due to finite sample effects. Closely related and with the same implications as sample overlap is *weak instrument bias*, which is generated by statistically weak SNP–exposure associations and related to the “*winner’s curse*” problem (Haycock et al., 2016). A recent preprint (Sadreev et al., 2021) examines the impact of weak instrument bias and winner’s curse in UK Biobank. Instrument strength is commonly measured via the F-statistic and a value of *F >* 10 is conventionally considered to guard against weak instrument bias (Burgess and Thompson, 2011; Davey Smith et al., 2020). In order to avoid bias from sample overlap, weak instruments and winner’s curse, a three-sample MR approach can be taken where, in addition to disjoint data for the SNP–exposure and SNP–outcome associations, the initial step of selecting genetic variants as instruments is based on a third dataset (Burgess et al., 2020a). However, three-sample MR analyses remain the exception.

Bias can also arise due to *reverse causation* if the effect of the genetic variant on the exposure is not primary (Burgess et al., 2021). In cases where a causal effect exists but the true causal direction between (hypothesised) exposure and (hypothesised) outcome is unknown, i.e., when it is not clear from background knowledge which variable is the cause and which variable the effect, then both SNP–exposure and SNP–outcome associations may reach genome-wide significance. Selecting the “wrong” variable as the exposure means the model is mis-specified, leading to erroneous effect estimates. Furthermore, time-dependent causal effects and feedback mechanisms can lead to causal links in the reverse direction. To avoid model misspecification, bi-directional MR (Timpson et al., 2011), which requires knowledge of valid instruments for both exposure and outcome, or Steiger filtering (Hemani et al., 2017) can be used as indicators for the correct causal direction.

In terms of the overall validity of genetic variants as instrumental variables, there exists potential bias due to *non-random inheritance* or assortative mating, giving rise to so-called “dynastic” effects. Within-family MR methods (Davies et al., 2019) have been proposed to adjust for mean parental genotypes. However, these biases predominantly affect socially influenced variables such as educational attainment and are rarely an issue in MR for many biological processes (Davey Smith et al., 2020; Brumpton et al., 2020).

Lastly, when possible, covariate-adjusted summary associations in MR should be avoided, as conditioning on a collider or on heritable covariates can bias GWAS outcomes (Hartwig et al., 2021). However, standard sources of confounding that pose major challenges for neuroimaging data (e.g., MRI artifacts, age, scanning parameters, etc.) should not be problematic in the context of MR analyses.

## 3 Neuroimaging data: Examples and applications

Any MR analysis should start with a hypothesis about a putative causal relationship between an exposure variable and an outcome variable. In neuroimaging, it may not be obvious *a priori* whether an image-derived phenotype (IDP) is the putative cause or the effect (or part of a feedback loop) in relation to other phenotypes. In some cases, a biologically plausible hypothesis for the true causal direction may be used as a basis for the MR model. For example, one may start by assuming that blood pressure causally affects certain IDPs, rather than variation in an IDP being a cause for changes in blood pressure levels; or one may assume that dMRI-derived connectivity features influence cognitive abilities, rather than the reverse being the case. However, without full knowledge of the biological pathways involved, bi-directional MR analyses (i.e., testing and comparing model fits for both causal directions) should be employed to guard against ill-specified models due to reverse causation.

From a modelling perspective, some common characteristics of neuroimaging data can exacerbate the challenges one may encounter in an MR analysis. First, one of the most prominent challenges of using Mendelian randomisation on neuroimaging data – now and for the foreseeable future – is sample size. MR requires huge sample sizes on the order of (at least) tens of thousands of individuals. This is only feasible with population-scale datasets. Even with datasets such as UK Biobank (*N ≈* 500k), the imaging cohort is typically much smaller (currently *N*_img_ *≈* 45k). Small sample size means reduced power to identify SNPs (via GWAS) as potential instrumental variables for MR, and consequently reduced power to detect a causal effect.

In MR analyses where IDPs are used as exposure, fewer (and weaker) instruments lead to bias towards the null in the MR estimate, as a consequence of regression dilution. If IDPs are used as outcome variables, higher variance in the SNP–outcome association leads to larger weights on individual SNP estimates in the IVW MR model and again a bias towards null. Higher variance in SNP–exposure and SNP–outcome associations will also increase confidence intervals of the final estimate.

A second challenge relates to weak instrument bias (possibly as a result of small sample size). Weak instrument bias can play a significant role in cases where IDPs are used as the exposure and only few, weakly associated SNPs are available as instruments. For instance, a recent study did not find any causal effects of IDPs on depression, but the authors note that a greater number of genome-wide significant SNPs associated with IDPs are needed before confident conclusions can be made (Shen et al., 2020).

A third challenge stems from the nature of brain phenotypes as “high-level” traits. In the biological causal chain, IDPs are far removed from the direct effects of genetic variation. Compared to “low-level” biomarkers (for example, proteins and their expression levels), SNPs are likely to be weaker instruments and more prone to pleiotropic bias when paired with biologically more distant phenotype exposures. Using IDPs (or other high-level phenotypes) as variables of interest in MR therefore means that the IV-assumptions are more likely to be violated. Conversely, a direct effect (short pathway) between SNP and exposure, and SNP and outcome generally reduces the possibilities of additional pleiotropic pathways. Neuroimaging MR analyses in particular are thus likely to require careful consideration of potential pleiotropic effects. Recent studies have relied on multi-variable MR approaches to account for (known) pleiotropy between multiple IDPs (Mo et al., 2021). Additionally, high-level phenotypes can be assumed to be more likely to exhibit non-linear associations. Non-linear MR approaches (Staley and Burgess, 2017) may therefore be particularly suited to causal investigations involving IDPs. Neuroimaging phenotypes may also be suitable candidates for an MR analysis based on polygenic risk scores (Dudbridge, 2021) as an alternative to multiple highly correlated IDPs.

Further considerations when planning an MR analysis on neuroimaging data may involve (i) heritability and (ii) unwanted confounding. Firstly, MR analyses using highly heritable phenotypes will have greater power to detect a causal effect. For example, white matter microstructure has higher SNP heritability (20–60%) compared with other neuroimaging modalities, indicating a greater genomic contribution to individual differences in phenotypes (Elliott et al., 2018). On the other hand, head movement and head size are usually adjusted for as nuisance covariates. Both are heritable attributes and known to be associated with certain personality traits. Conditioning on any nuisance covariate that is associated with the outcome and the instrumental variables, and/or the exposure, can introduce a spurious association between SNPs and outcome (collider bias, (Munafò et al., 2018). Therefore, deconfounding of certain imaging confounds can lead to adding rather than eliminating sources of bias in the context of MR.

Confounds are a major nuisance in neuroimaging applications and include motion artefacts, age, scanner- and site-specific factors, head size and various other potential confounding variables. In theory, assuming the IV-assumptions hold and the SNPs used for analysis are valid instruments, MR estimates are not biased by confounders of the exposure–outcome relationship, thereby removing the necessity to deconfound neuroimaging phenotypes prior to analysis. In practice, GWAS association estimates involving IDPs are routinely deconfounded for large sets of covariates (Alfaro-Almagro et al., 2021). This can introduce bias in MR outcomes if the covariate is a collider (common effect) on the pathway linking the SNP to the IDP of interest (Hartwig et al., 2021). The most likely source of confounding, however, stems from population stratification. A recent study on UK Biobank data has shown that selection bias may be compounded in the case of imaging phenotypes compared to other variables (Lyall et al., 2021). And since only a few population-wide (and openly available) datasets exist, any inherent bias is more likely to influence results across multiple studies, as independent research groups rely on the same data for their analyses.

Finally, the potential impact of time-dependent effects should be taken into account in neuroimaging applications. Feedback cycles and time-varying exposures (Shi et al., 2022) are commonly ignored in MR methods, but may have a non-negligible influence in certain imaging contexts. For example, longitudinal imaging data would be necessary to determine the interplay of neurodevelopmental and neurodegenerative factors of a disease mechanism. In aetiological disease research, MR can help to investigate differences between environmental and genetic disease risks (Storm et al., 2020). While one cannot expect that estimated MR effect sizes will predict the effect size of a medical intervention (gene–environment non-equivalence) – since the SNPs represent a life-time exposure to a weak version of the risk factor – MR can still be used to discover risk factors that are potential drug targets for drug development, without the limitations of observational studies and the implications of carrying out RCTs. However, one should be aware that MR effect size estimates are unlikely to correspond to effect magnitudes of a medical intervention, and the pathways involved are almost certainly different.

In the following sections, we look at three different real-data applications (selected via a wide-ranging exploratory analysis) in which neuroimaging features (IDPs) play a role as exposure or outcome in Mendelian randomisation analyses.

### 3.1 Data

We use the recently expanded UK Biobank GWAS database (Smith et al., 2021) of summary statistics for over 17 million SNPs, to identify associations between genetic variants and brain imaging derived phenotypes. The current release of multimodal imaging data comprises almost 4,000 individual measures (IDPs) from over 33,000 participants. The full UKB cohort for which non-imaging data is available has a sample size of about 500,000 (Miller et al., 2016). Additionally, we use openly available GWAS results from large international consortia via the MRC IEU OpenGWAS infrastructure (Elsworth et al., 2020) when using two-sample MR to identify SNPs associated with non-imaging derived phenotypes (nIDPs). Each GWAS controlled for different nuisance variables, which typically includes sex, age, and the main components of genetic population variation. Additionally, all IDP data had extensive nuisance modelling as described in (Alfaro-Almagro et al., 2021).

#### 3.1.1 Pre-selection of potentially causal associations

Due to the large number of variables available in UK Biobank (UKB), we employed a pre-selection procedure in order to reduce the number of exposure–outcome pairs involving 3,935 IDPs and 4,178 nIDPs. Although not every possible combination is biologically plausible, for the vast majority of imaging-related traits, causal links to other phenotypes and health outcomes are not yet established in the scientific literature. We therefore took an *a priori* agnostic approach to variable selection.

Based on heritability estimates via LD score regression (Bulik-Sullivan et al., 2015) for each nIDP (data made available by the Neale lab), we set the cut-off for the heritability significance level at *p <* 0.05 and required a confidence metric^8^ of medium or higher. Additionally, we set a maximum allowed value for the LD score intercept of 1.1, as a larger intercept value may indicate population stratification, confounding or other sources of model misspecification. This resulted in 922 highly heritable nIDPs with heritability estimates in the range of 0.05–0.40. A schematic description of the selection procedure in form of a flowchart is given in Figure 12.

IDPs were filtered analogously using heritability data published alongside GWAS results via Oxford’s BIG 40 Brain Imaging Genetics Server (Smith et al., 2021). Due to generally high heritability of brain phenotypes, the majority of IDPs (n=2706) were retained.

For a further selection step, we computed pairwise correlations between all IDP–nIDP pairs that survived the heritability selection. Filtering the correlation results, we set a threshold for absolute correlation values at *|ρ| >* 0.10 and a minimum significance level of *−* log_10_(*p*) *>* 12 (noting that the Bonferroni threshold would be 7.7), resulting in 895 correlated IDP–nIDP pairs, with 365 unique IDPs and 133 unique nIDPs.

In this unbiased approach we are not pre-specifying which variables are outcomes and which are exposures. Typically, MR analyses start with a specific exposure and outcome of interest, often informed by background knowledge on specific biological pathways. Here we adopt a more exploratory approach. The reasons for this are two-fold: First, the causal mechanisms involving imaging phenotypes are generally not well understood and thus there is limited prior information available on which to base a well-formed hypothesis that could be tested via MR. Second, we strongly emphasise the need for a rigorous sensitivity analysis (see Table 1) to reduce the possibility of the results being susceptible to the problem of reverse causation and other biases.

For illustrative purposes (see third example in Section 3.3), we also handpicked a set of seven nIDPs related to cognition in UK Biobank. These include “duration to complete alphanumeric path”, “duration to complete numeric path”, “fluid intelligence score”, “maximum digits remembered correctly,” “mean time to correctly identify matches”, “number of puzzles correctly solved” and “number of symbol digit matches made correctly.” Here, we set the thresholds for correlations with IDPs at *|ρ| >* 0.05 and *−* log_10_(*p*) *>* 10.

#### 3.1.2 Screening of potentially causal associations

Next we carried out a preliminary screening for causal effects by running a standard IVW MR analysis on each of the 895 previously identified potential causal pairs, considering each variable as exposure and outcome in turn, using the TwoSampleMR R package (Hemani et al., 2018b). For each variable, we used GWAS on European ancestry subjects with largest sample sizes from the MRC IEU GWAS database to identify SNPs that are strongly associated with any of the nIDPs. In many cases this meant that UKB GWAS results were chosen for both IDP and nIDP variables, thereby creating a sample overlap between exposure and outcome GWAS summary statistics. Because the imaging cohort and thus sample size for IDP GWAS is much smaller than the total UKB sample (to date, around 10% of UKB participants have been imaged), this can still be considered a two-sample MR setup. Potential issues due to sample overlap are expected to be small and may result in reduced sensitivity to detect an effect (see Section 2.4). Genetic variants were harmonised using default parameters in the TwoSampleMR package.

We note that screening approaches, like the one we have used here, should only be considered in exploratory settings. Related to issues arising from multiple comparisons, there is a danger that results are selected that are based on inflated associations due to random noise by chance.

Sufficiently strongly associated SNPs could not be identified for some of the 895 variable pairs retained in the pre-selection step, thereby reducing the number of available IDP–nIDP pairs to 620. The screening with IVW MR then resulted in a total of 1240 bi-directional causal estimates for 620 variable pairs. We selected three example scenarios involving blood pressure, bone density and cognition, respectively, and their associations with various IDPs. These sets of variables are among the strongest effects observed in the screening phase.

Overall, we found 449 significant effects (*p <* 0.05), 32 of which involved an IDP as exposure, an nIDP as outcome and an average of 25 genome-wide significant SNPs as instruments (range 2–45). The majority of exposure IDPs were volume-based measures (21), the rest were diffusion-derived IDPs (10) and one surface measure. The remaining 417 significant effects involved an nIDP as exposure (with the largest group of 153 based on blood pressure) and an IDP as outcome, with an average of 303 genome-wide significant SNPs (range 4–731) per MR analysis. The majority of outcome IDPs were based on diffusion-derived metrics (267), with the rest including volume (121), intensity (24) and surface measures (5). Detailed results of all MR screening analyses are included in the Supplement (Supplementary File 1).

The example selection is also motivated by our intention to showcase different application scenarios and highlight potential challenges, together with possible approaches of how to deal with these, involving existing software tools and statistical checks. The examples include structural as well as diffusion-based IDPs, and show cases in which IDPs are hypothesised as being affected by biophysical phenotypes (blood pressure and bone density) and one case in which IDPs may be assumed as a putative cause of (small) differences in cognitive ability. The first example shows a strong, statistically robust effect, whereas the other two are less clear-cut, warranting careful and detailed sensitivity analyses.

### 3.2 Methods

For each of the three selected example scenarios, we performed a detailed MR and sensitivity analysis. Additionally, we fitted Bayesian networks for the first example, including a set of covariates which could be expected to act as confounders of the exposure and outcome association.

Apart from the standard IVW MR approach, we included weighted median- and mode-based MR as well as meta-analytic IVW variants with fixed and multiplicative random effects. The (multiplicative) random-effects model allows for over-dispersion in the regression model and therefore can account for some heterogeneity in the causal estimates of individual SNPs (Burgess and Bowden, 2015). Robust methods included MR-Egger, MR-RAPS, MR-Mix and MR-PRESSO.

The sensitivity analysis included the Steiger directionality test, outlier detection and removal via MR-PRESSO and heterogeneity tests via Cochran’s *Q* statistic for several MR methods. We also varied the SNP–exposure association threshold for instrument selection. In addition to the commonly used default of *p <* 5 *×* 10*^−^*^8^, the MR analysis was repeated with a more liberal (*p <* 10*^−^*^6^) and a more conservative (*p <* 10*^−^*^12^) selection threshold. In cases where only weak associations with the exposure of interest could be identified from GWAS data (mostly affecting IDPs), the stringent and default threshold options were omitted as they would result in an empty selection set, and only the liberal threshold was used.

Estimation of Bayesian networks and the corresponding graphical structures was carried out with the bnlearn R package (Scutari, 2010). Specifically, we used the hill-climbing algorithm with BIC (Scutari and Denis, 2021), which includes necessary distributional assumptions about continuous (normally distributed) and discrete (multinomially distributed) variables in the network. Bootstrapping of the data resulted in 1000 replications per setting, which were used to estimate the likelihood of edge inclusion, following a similar setup as in Howey et al. (2020). The probability of an edge existing, and the probability of the edge being in a particular direction (given that it exists) were estimated by counting the proportion of times that such events occurred amongst the 1000 resulting best-fit bootstrap networks.

### 3.3 Results

#### 3.3.1 Example 1: Blood pressure

The strongest MR effect estimates based on the screening of candidate exposure–outcome pairs resulted from using blood pressure related measures as exposure. Modelling blood pressure as the exposure rather than the outcome in an MR analysis can reasonably be motivated using biological arguments. In order to investigate the potential effects of blood pressure on different parts of the brain, we focused in the detailed analysis on the following three IDPs that showed some of the strongest IVW MR effects: i) volume of white matter hyperintensities (WMH) measured on T2 FLAIR images, ii) mean diffusivity in the superior longitudinal fasciculus, and iii) mean diffusivity in the external capsule. Bi-directional effect estimates are shown in Figure 4 using 396 SNPs for the forward analysis (panel A) and 7 SNPs for the reverse direction (panel B). The reverse direction results are not significant, having confidence intervals that mostly cover zero.^9^

**Figure 4:**
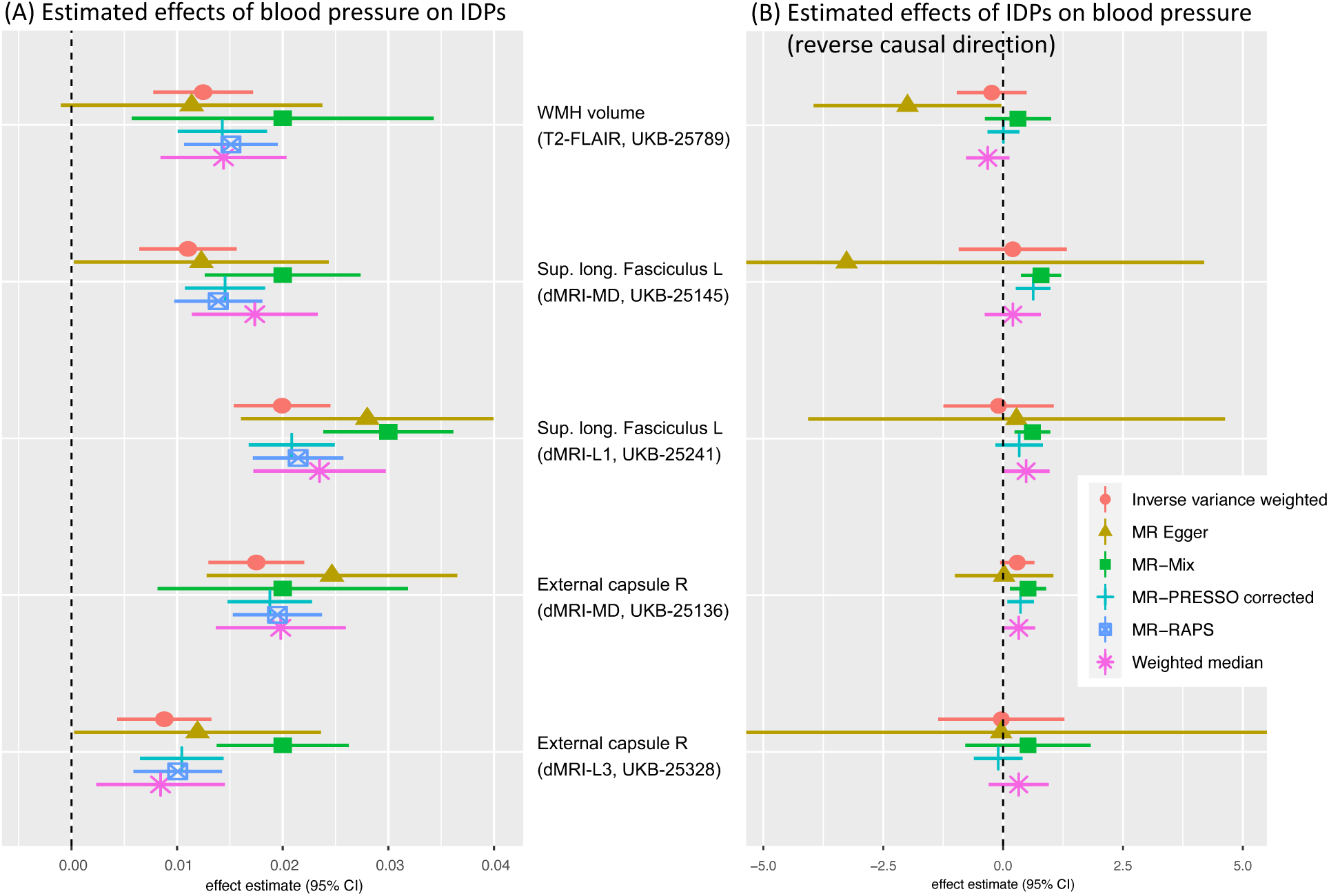
Bi-directional MR analysis of the relationship between systolic blood pressure and selected IDPs: Shown are causal effect estimates for six MR methods. (A) Causal effect estimates of BP on IDPs. (B) Causal effect estimates of IDPs on BP. Errorbars show 95% confidence intervals.

**Figure 5:**
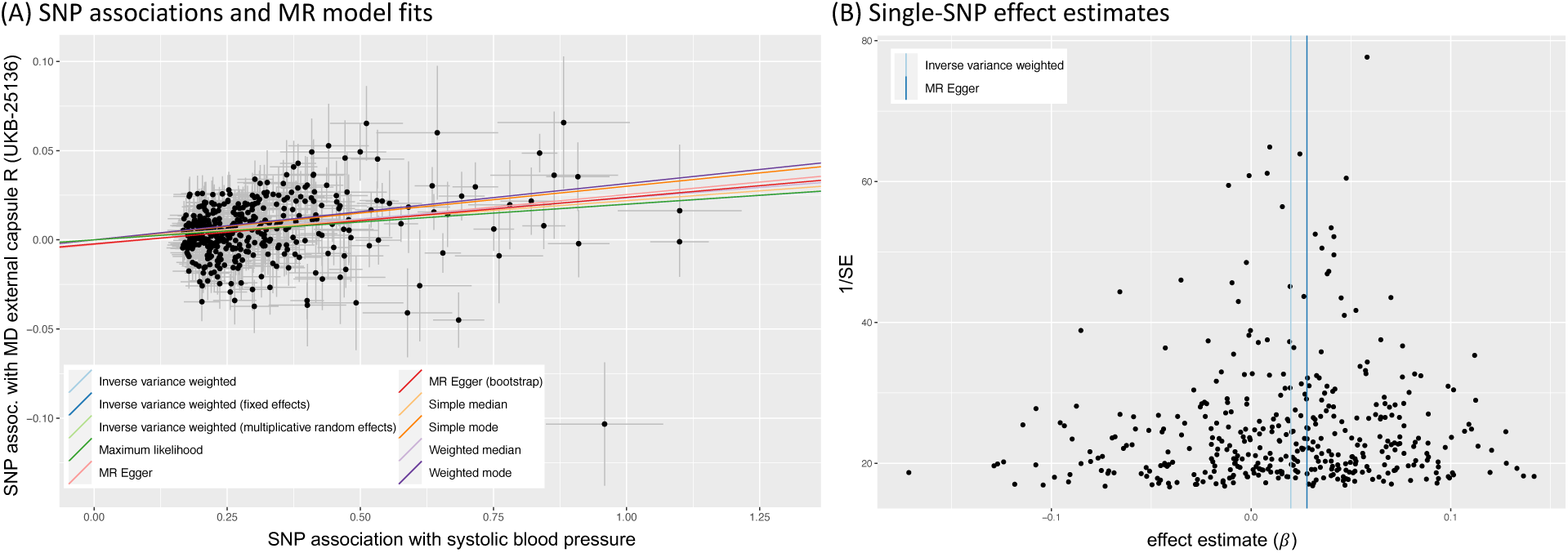
MR sensitivity analysis of the causal effect of systolic blood pressure (exposure) on mean diffusivity of the external capsule WM tract R (UKB-ID 25316). (A) Scatter plot showing associations of individual SNPs with exposure and outcome variables, with dotted cross-hairs indicating standard errors. Coloured lines indicate effect estimates from regression fits using different MR methods. (B) Funnel plot of single-SNP effect estimates and corresponding inverse standard errors.

To illustrate the steps laid out in Table 1, Figures 5 and 13 show several plots from a sensitivity analysis for the effect of systolic blood pressure (BP) on the mean diffusivity (MD) in the right external capsule as measured by diffusion MRI (UKB-ID 25136). The forward-direction results in Figure 4 suggest a causal effect of BP on these IDPs of 0.01 to 0.03; for example, for every SD difference in BP, approximately a 0.02 SD difference in external capsule MD is expected. Visual inspection of the scatter plot in Figure 5A reveals a consistently strong positive causal effect of BP on the right external capsule.

The MR-Egger intercept was not significantly different from zero (*p* = 0.1), indicating a lack of evidence for horizontal pleiotropy and supporting the exclusion assumption that the only pathway from selected SNPs to the outcome is via the exposure. Statistical heterogeneity tests using Cochran’s *Q* statistic (*Q* = 596*, df* = 395*, p* = 2 *×* 10*^−^*^10^) indicate substantial heterogeneity in the individual effect estimates. However, as can be seen in the funnel plot in Figure 5B, there is no strong pattern of asymmetry and therefore no clear indication of unbalanced, directional pleiotropy that could bias the final effect estimate. The non-significant MR-Egger intercept together with the approximately symmetric distribution of individual effects in the funnel plot may indicate that, overall, pleiotropic effects balance out and thus are unlikely to invalidate the MR result.

The MR-PRESSO outlier test found three potentially problematic SNPs. Removing these three SNPs resulted in a similar causal effect estimate (*β* = 0.02) and a higher significance level (*p* = 5 *×* 10*^−^*^21^ vs. *p*_uncorr._ = 5 *×* 10*^−^*^18^). Single-SNP and leave-one-out analysis (Figure 13) also do not show substantial outliers that would indicate that the effect estimate is driven by any single SNP.

A Steiger directionality test (*p* = 0.0049) indicated that the more likely causal direction is such that BP affects the external capsule and not the other way around. As can be seen in the effect estimates for the reverse causal direction (Figure 4B, *n*_SNP_ = 7), there is no indication for a causal pathway originating with an IDP and causing changes in BP. Unfortunately, due to the much weaker SNP–IDP association strengths (partly due to lower sample size), it is difficult to ascertain whether this reflects a truly uni-directional influence of BP or simply insufficient power to detect a causal influence in the reverse direction.

Performing the same MR analysis with a stricter SNP–exposure association threshold reduced the number of instruments from 396 to 219, with similar causal effect estimates for all MR methods and slightly increased corresponding p-values.

Taken together, the results from robust MR methods and sensitivity analyses described above do not indicate the presence of strong pleiotropic effects or other sources of systematic bias that could substantially influence the MR results. Consistent results from standard, robust and outlier-removal MR variants, combined with the absence of directional pleiotropy as assessed by the MR-Egger intercept and the meta-analytic funnel plot, strongly support a causal effect of systolic blood pressure on the external capsule. Furthermore, there is no evidence for reverse causation, which is in line with what could reasonably be expected in terms of biological pathways.

We have not explicitly considered the potential for correlated pleiotropy. In case of the current application, a hypothesised correlated pleiotropic pathway could include BMI, which is known to be an important determinant of blood pressure. If BMI also has an effect on the outcome, it could bias the MR estimate but might not be easily detected if more than a few outlying SNPs are affected. This could be addressed through multivariable MR or via an approach that is robust to correlated pleiotropy such as MR-CAUSE, which has been shown to reduce false positives when correlated pleiotropic pathways are present (Morrison et al., 2020).

In addition, a Bayesian network analysis can be used to corroborate findings from the MR analysis or, conversely, MR results can provide prior knowledge about (i.e., constraints on) the presence or absence of directed edges in the underlying graph structure. In this example, the set of variables to estimate the graph structure included systolic blood pressure (BP) and MD in the external capsule (EC) as primary variables of interest, as well as additional covariates Age, Grey Matter volume (GM) and Income (selected to reflect associations with socio-economic background). Five SNPs that are strongly associated with either BP, GM or EC were included as directional anchors to increase the identifiability of edges in the graph.

The Bayesian network shown in Figure 6A was estimated using the hill-climbing algorithm in the bnlearn R package and is based on 1000 bootstrap samples. A threshold of 0.8 for edge-inclusion was used, i.e., requiring that a candidate edge is present in at least 80% of bootstrap samples. A comparison (not shown) between the bootstrapped graph and a single estimation on the full data was used to highlight any discrepancies or inconsistencies (none were found in this case). Using directional anchors (indicated as solid arrows in Figure 6A), the BN estimate results in a DAG with a strong edge from BP to the external capsule. Similarly the ancestral graph estimate (Figure 6B) confirms the expected causal direction from BP to EC, albeit with the caveat that one or more latent, unmeasured variables may be present in the path from BP to EC.

**Figure 6:**
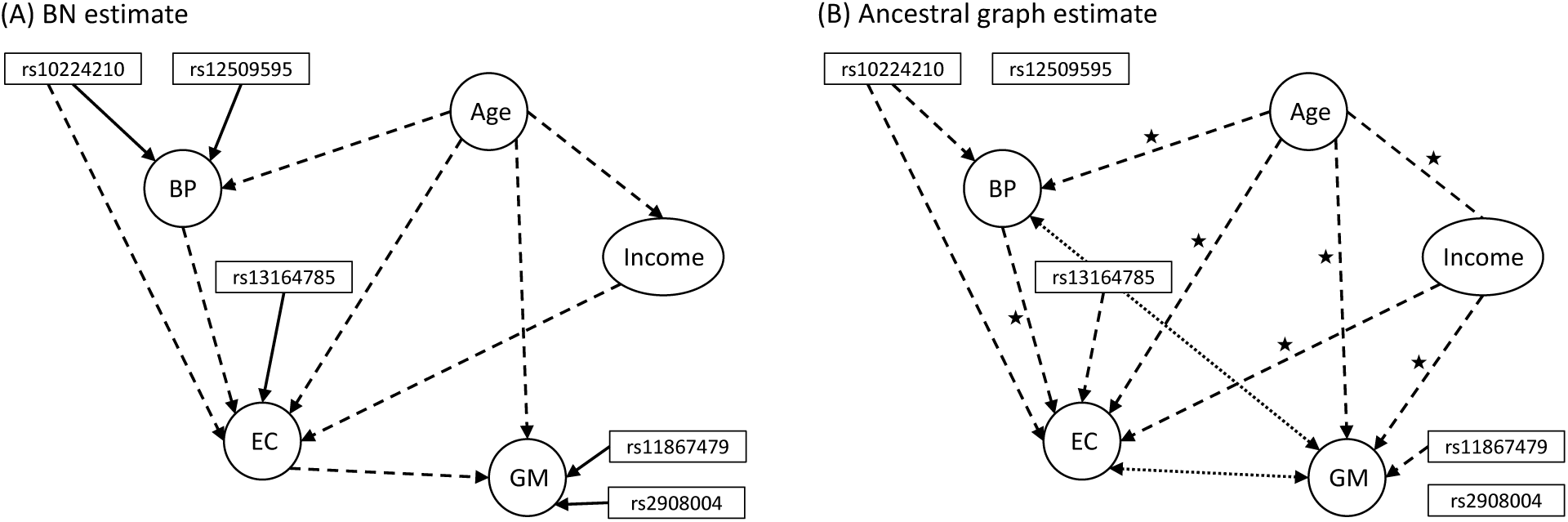
Bayesian network estimates involving systolic blood pressure (BP), mean diffusivity of the external capsule WM tract (EC) and three covariates (Age, Income, grey matter (GM) volume). Estimated edges are denoted with dashed arrows. (A) Standard BN estimate using the hill-climbing algorithm with BIC in bnlearn. Five SNPs (with known associations with the connected phenotype) are used as genetic “anchors”, i.e., the known associations are provided as prior information and are represented as fixed edges in the graph (indicated as solid arrows). Edges between SNPs and those that would have Age or any SNP as effect (endpoint of an arrow) were excluded *a priori*. (B) Ancestral graph estimate based on the RFCI algorithm in the pcalg R package; no prior information on presence or absence of individual edges was provided. No edges were found for two SNPs. Dotted, bi-directional arrows indicate the presence of a common cause. An edge without arrowheads means that directionality of the relationship could not be determined from the data (e.g., Age–Income). A ⋆ next to an edge indicates the potential presence of a latent, unmeasured variable.

#### 3.3.2 Example 2: Bone mineral density

The second example looks at bi-directional effects involving heel bone mineral density (UKB-ID 3148) and various IDPs. A priori, one would not necessarily expect brain phenotypes to have a causal influence on bone density. However, the reverse direction is not biologically obvious either, and one might hypothesise that indirect effects may play a role. We set the default “forward” direction of any causal relationship as going from bone density to IDP. Results from a bi-directional MR investigation are shown in Figure 7, highlighting six IDPs with the strongest overall effect estimates.

**Figure 7:**
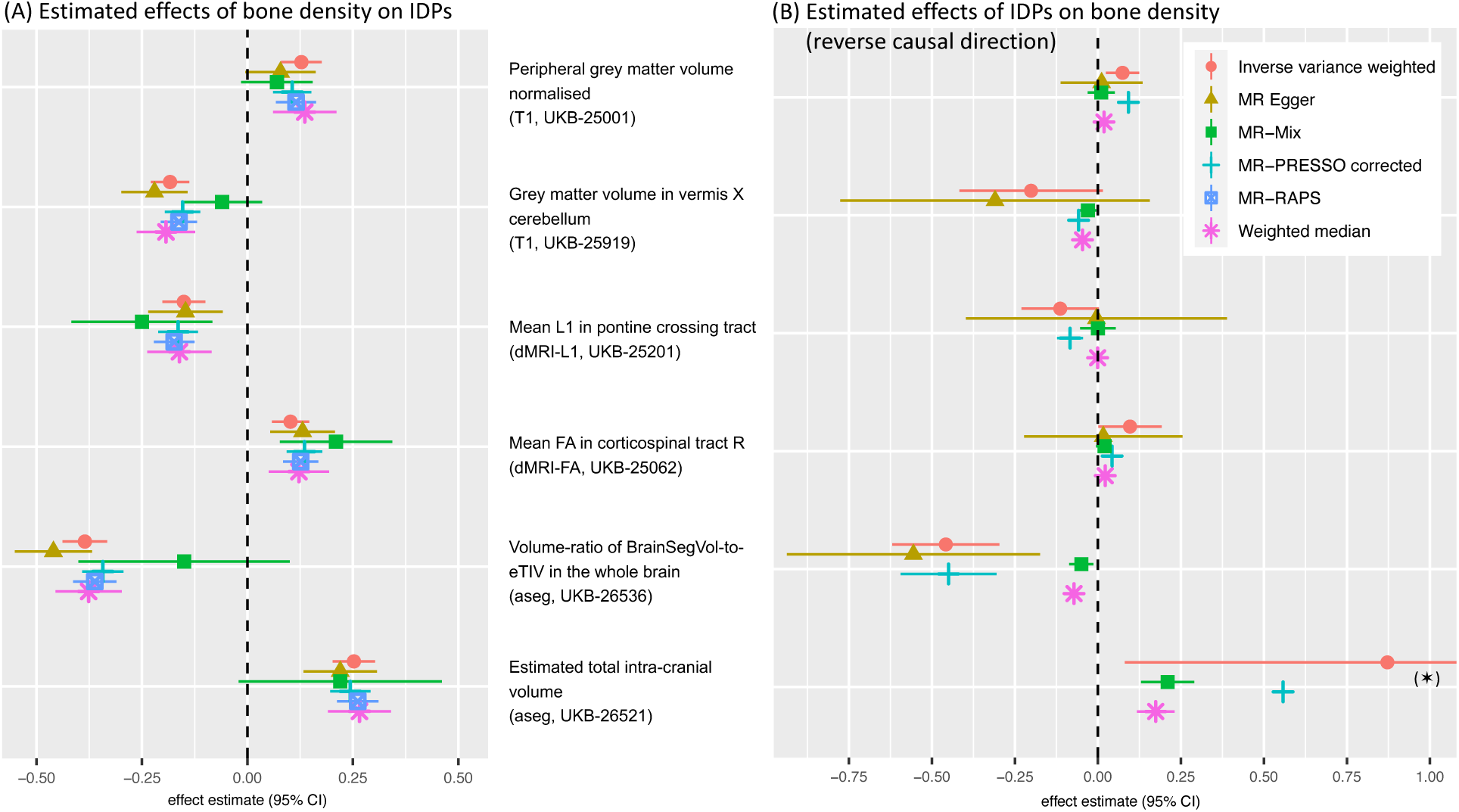
Bi-directional MR analysis of the relationship between heel bone mineral density (UKB-ID 3418) and selected IDPs: Shown are causal effect estimates for six MR methods. (A) Causal effect estimates of heel bone mineral density on IDPs. (B) Causal effect estimates of IDPs on heel bone mineral density. Errorbars show 95% confidence intervals. (⋆) MR-Egger estimate (2.9 *±* 0.8) not in plotting range.

Figure 7A shows positive and negative effect estimates of bone density on both structural and diffusion-based IDPs. On the other hand, there is no indication for reverse causation for most IDPs except for two Freesurfer measures related to brain volume.

For the remainder of this example, we focus on T1 normalised peripheral cortical grey matter (GM) volume generated via FSL (UKB-ID 25001) and show results from a sensitivity analysis. The scatter plot of SNP associations for the forward direction (Figure 8A) shows much more consistent effect estimates across different MR methods than the same plot for the reverse direction (Figure 9A). Substantial heterogeneity is detected by Cochran’s *Q* test in both directions, although it is more severe for the reverse effect estimates (*Q* = 400*, df* = 30, *p <* 10*^−^*^66^) than for the forward direction (*Q* = 734*, df* = 433, *p <* 10*^−^*^18^). Strong heterogeneous and asymmetric effects can also be seen in the funnel plot in Figure 9B.

**Figure 8:**
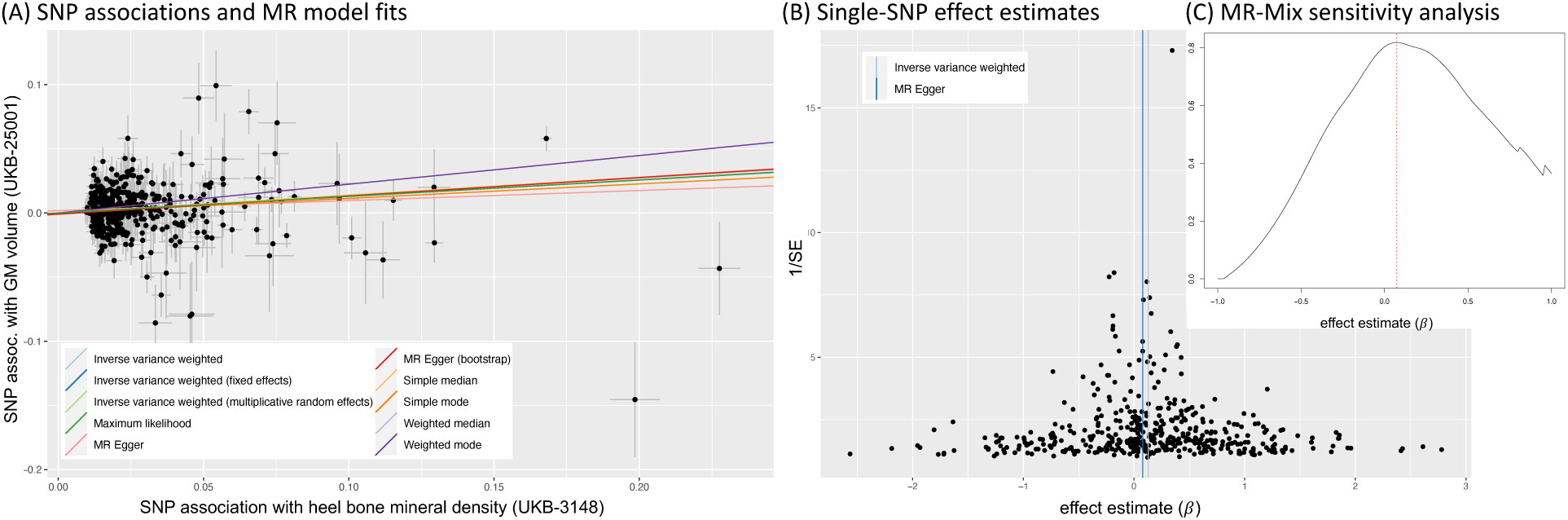
MR sensitivity analysis of a potentially causal effect of heel bone mineral density (UKB-ID 3148) on T1 peripheral cortical GM volume (UKB-ID 25001). (A) Scatter plot showing associations of individual SNPs with exposure and outcome variables. Coloured lines indicate effect estimates from regression fits using different MR methods. (B) Funnel plot of single-SNP effect estimates and corresponding inverse standard errors. (C) Probability density of the estimated causal effect using a mixture model approach with MR-Mix. The dashed red line indicates the causal effect estimate.

**Figure 9:**
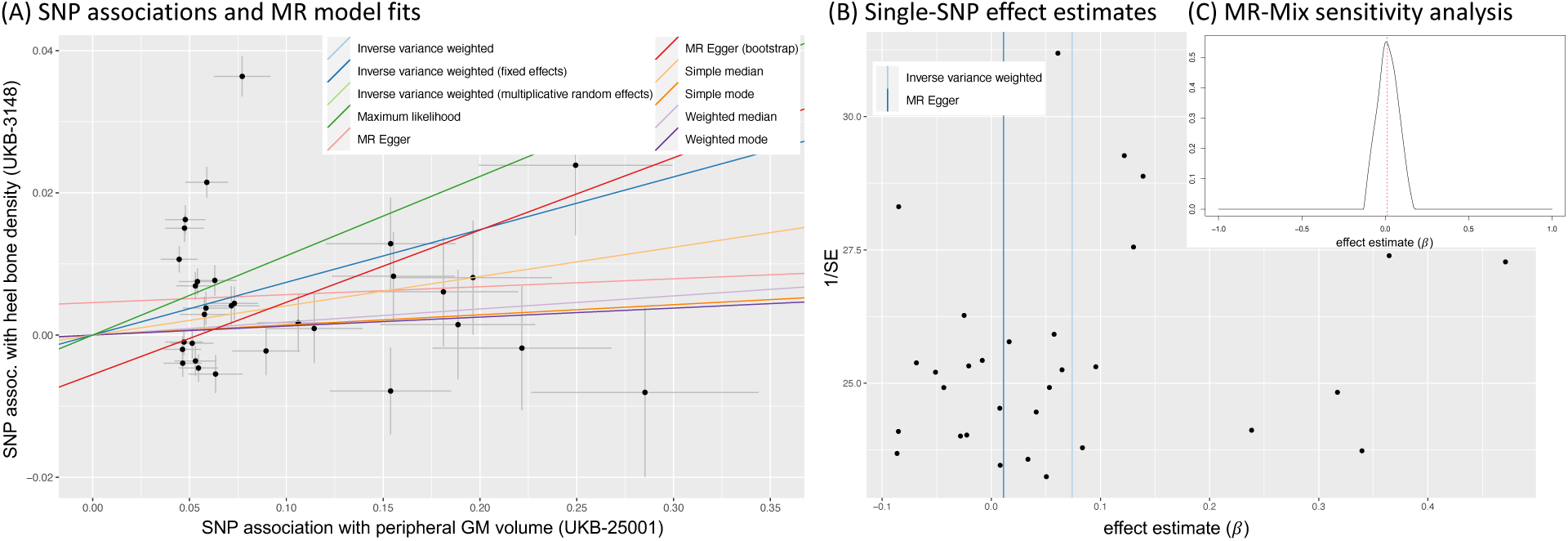
MR sensitivity analysis of the effect of T1 peripheral cortical GM volume (UKB-ID 25001) on heel bone mineral density (UKB-ID 3418). (A) Scatter plot showing associations of individual SNPs with exposure and outcome variables. Coloured lines indicate effect estimates from regression fits using different MR methods. (B) Funnel plot of single-SNP effect estimates and corresponding inverse standard errors. (C) Probability density of the estimated causal effect using a mixture model approach with MR-Mix. The dashed red line indicates the causal effect estimate.

Scrutiny of the MR-Mix (Qi and Chatterjee, 2019) output demonstrates the discrepancy in effect estimates. The density fit in Figure 9C for the reverse direction is much narrower and suggests a high confidence that the effect is close to zero.^10^ Furthermore, the MR-PRESSO fit flags 24 out of 31 SNPs in the reverse analysis as potential outliers, and a Steiger directionality test indicates the forward direction as the more plausible pathway.

Overall, evidence from this analysis points more strongly towards a forward causal effect of bone mineral density on GM volume rather than the reverse. However, feedback mechanisms cannot be fully ruled out and the potential existence of a common cause that is also associated with the selected instruments (violation of the exclusion restriction) could severely bias the MR results.

In cases like this, domain knowledge and other sources of evidence are crucial and will increase confidence in any findings from an MR analysis and help in the interpretation of results.

#### 3.3.3 Example 3: Cognition

For this example, we use a set of variables related to cognitive function to demonstrate some of the challenges that may arise in the study of causal relationships of brain phenotypes, especially when using IDPs as exposures in an MR framework. During the preliminary screening of potential causal variable pairs, a reaction time measure, “mean time to correctly identify matches” (MTCIM, UKB-ID 20023), showed the strongest associations with IDPs. Figure 10 gives causal effect estimates with MTCIM as (A) the exposure and (B) the outcome variable, respectively. Unlike in the previous two examples, effect estimates are more variable and generally closer to zero, meaning that statistical significance as measured by p-values is lower. Lower levels of significance and higher variability can in part be attributed to fewer and weaker instruments being available for a high-level trait such as cognition, compared to phenotypes such as blood pressure, for example, which are arguably biologically closer to the direct effect of genetic variants. For the vast majority of IDPs, and based on currently available GWAS data, only a handful of SNPs are strongly associated with an IDP in any given case, thus limiting the power to detect true causal effects.

**Figure 10:**
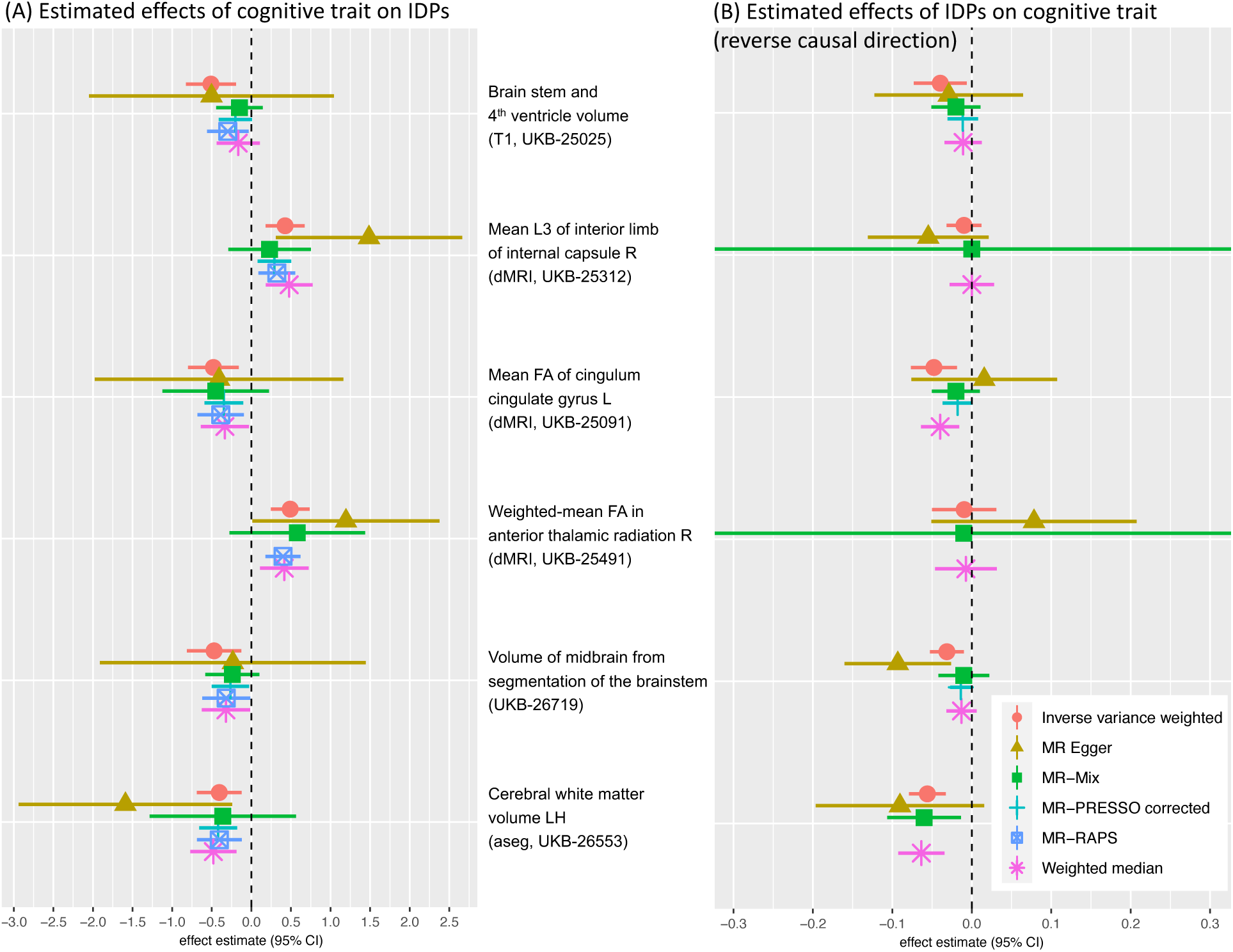
Bi-directional MR analysis of the relationship between the mean time to correctly identifying matches in a cognitive test (UKB-ID 20023) and selected IDPs: Shown are causal effect estimates for six MR methods. (A) Causal effect estimates of the cognitive trait on IDPs. (B) Causal effect estimates of IDPs on the cognitive trait. Errorbars show 95% confidence intervals.

Looking at one IDP (cerebral white matter volume in the left hemisphere, UKB-ID 26553) in greater detail, we performed a similar sensitivity analysis as in previous examples. The plots in Figure 10 show more reliable estimates for the forward direction (MTCIM effect on WM volume, *n*_SNP_ = 62) than for the reverse direction (*n*_SNP_ = 13). The funnel plots in Figure 11B,D show asymmetry in the single-SNP effect estimates for both directions, indicating the presence of directional pleiotropy.

**Figure 11:**
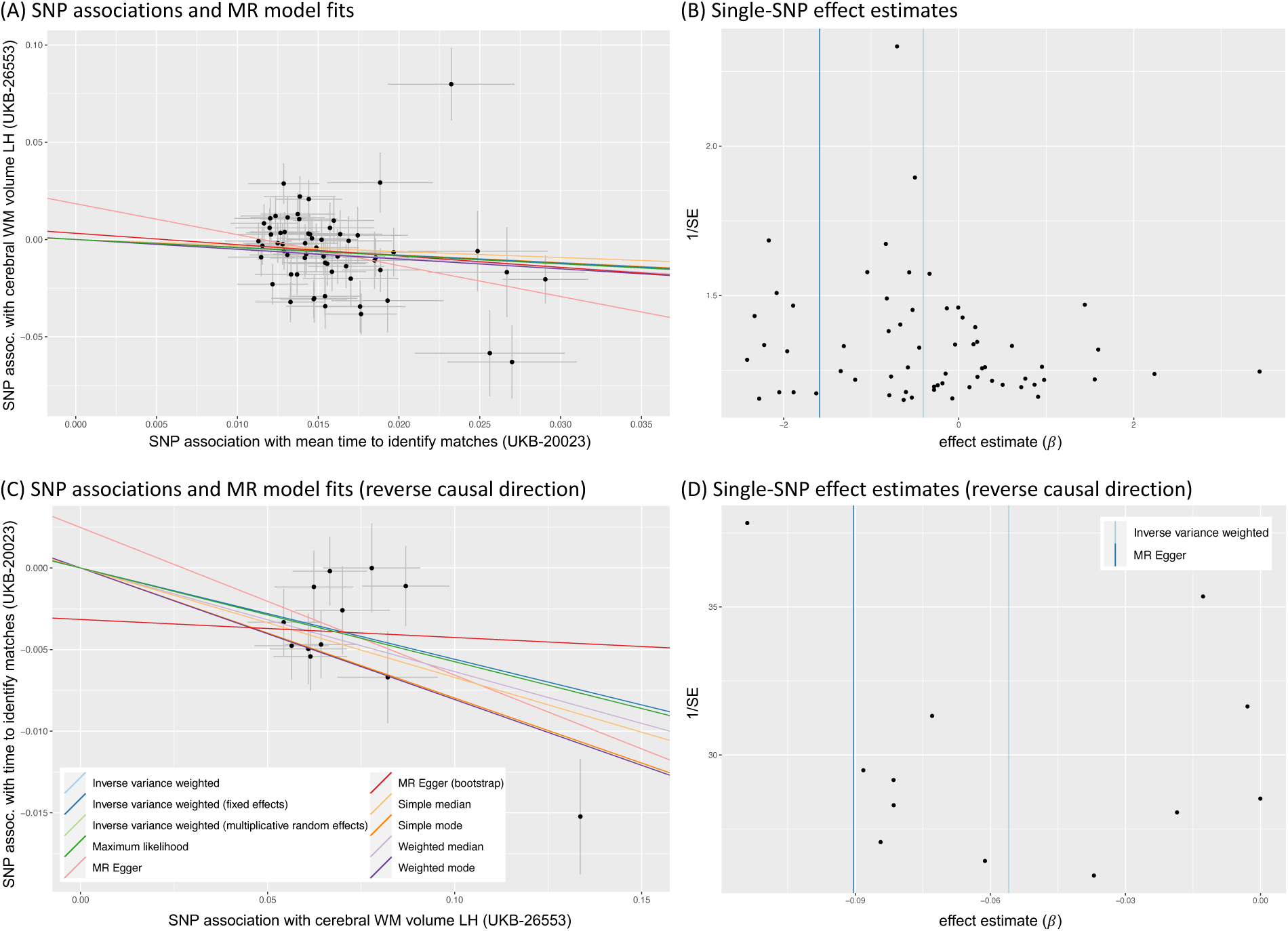
Bi-directional MR sensitivity analysis of the potentially causal effect of correctly identifying matches (UKB-ID 20023) on subcortical white matter volume (UKB-ID 26553) (A-B) and vice versa (C-D). (A) and (C) Scatter plots showing associations of individual SNPs with exposure and outcome variables. Coloured lines indicate effect estimates from regression fits using different MR methods. (B) and (D) Funnel plots of single-SNP effect estimates and corresponding inverse standard errors.

Cochran’s *Q* test showed greater heterogeneity in single-SNP estimates in the forward direction and the Steiger directionality test was inconclusive, i.e., estimating both directions as roughly equally likely based on the strength of the SNP–exposure and SNP–outcome associations. Single-SNP (Figure 17A) and leave-one-out MR analysis (Figure 17B) did not reveal any clear outliers.

A possible reason for the ambiguous bi-directional MR results could lie in an upstream pleiotropic phenotype that is a common cause for both exposure and outcome (correlated horizontal pleiotropy). To give one potential example, educational attainment could be hypothesised to affect each of the phenotypes considered in the MR, and thus bias the causal effect estimate even though no clear outliers have been detected.

Perhaps counter-intuitively, a potential causal effect from MTCIM to WM volume appears more plausible based on these findings. However, it is not clear what biological pathways may be involved. Additionally, feedback loops and time-dependent mechanisms (Burgess et al., 2021) may also play a role here. Overall, without further corroborating evidence, clear causal conclusions cannot be drawn.

We urge caution when interpreting and reporting results from potentially under-powered MR analyses or when a thorough sensitivity analysis indicates underlying issues with outliers, heterogeneity and pleiotropy. The burden of establishing credible evidence is particularly high in cases where existing domain knowledge is limited, and claims of newly discovered causal mechanisms are made.

## 4 Discussion

The goal of this paper was to introduce Mendelian randomisation to researchers with a predominantly neuroimaging background, and illustrate the advantages and limitations of MR with three neuroimaging-specific example applications. Although often not stated explicitly, causal claims about biological mechanisms and disease pathways are commonly made implicitly. MR methods can provide a framework to rigorously test causal hypotheses in the absence of interventional data, but should not be considered as the only source of evidence (and are not as bullet-proof as a randomised, controlled, interventional study). Additional domain knowledge and the incorporation of results from different methodologies are considered key ingredients for a *triangulation of evidence* approach (Howey et al., 2020; Lawlor et al., 2016; Munafò and Davey Smith, 2018). We briefly explored one such complementary approach in the form of Bayesian networks here. Causal Bayesian networks can be used as hypothesis-generating approaches that result in testable predictions about effects under an external manipulation of a parent node on its descendants (Grosse-Wentrup et al., 2016). These dependencies can in turn be corroborated or informed by findings from MR analyses.

Recent recommendations for best practices in MR include prioritising investigations in which the associations between genetic variants and exposures of interest are both primary (e.g., direct SNP effect on protein level) and well-understood (Burgess et al., 2021). In the field of neuroimaging this is generally not the case. Nevertheless, Mendelian randomisation can be used as a powerful tool to advance our knowledge of existing causal pathways and discover new ones. We strongly believe that thorough consideration of underlying assumptions, general limitations of MR, potential sources of bias alongside rigorous sensitivity analyses and cautious interpretation of results are necessary components of a careful MR investigation (see Table 1).

To our knowledge this is the first MR study involving hundreds of imaging-derived phenotypes together with a wide range of health measures. The purpose of this exploratory investigation was to showcase the potential of MR analyses when applied to neuroimaging data. We highlighted three exposure–outcome scenarios with the intention to demonstrate some of the methodological limitations and inherent difficulties with ensuring reliable results.

Our first example involved systolic blood pressure and its effect on various IDPs. Particularly, we found evidence for a robust causal effect of blood pressure on mean diffusivity in the external capsule. Results from sensitivity analyses corroborated this finding. We also compared MR results to graphical estimates using Bayesian networks in a complementary analysis.

In the second example, we focused on bi-directional effects involving bone density, either as possible exposure or possible outcome in an MR analysis. In the absence of clear, biologically grounded hypotheses about cause–effect directionality and robust one-directional effect estimates, MR results can be inconclusive and extra care about potential violations of underlying assumptions need to be taken. We showed in an extensive sensitivity analysis how one might use available software tools, statistical tests and plotting of the data to further investigate MR findings.

The third example was motivated by the possibility of (causally) relating imaging-derived phenotypes to high-level traits such as cognition. Highly variable effect estimates and low levels of statistical significance revealed substantial challenges when investigating traits that are far removed from the direct effects of genetic variation. Unfortunately, due to the much weaker SNP–IDP association strengths (which are partly due to comparatively small sample size of the imaging GWAS), most causal effect estimates involving IDPs, either as exposure or as outcome, are small and often inconclusive. Multi-variable MR methods, possibly combined with a dimensionality-reduction and orthogonalisation step for highly-correlated imaging exposures as recently proposed (Mo et al., 2021), are one direction of ongoing methodological development to address some of these issues.

We emphasise that, apart from the first example, which showed a robust and reliable causal effect of blood pressure on a measure of diffusivity in the right external capsule, our findings revealed only putative causal effects with indications of potential bias due to pleiotropy, weak instruments or reverse causation. Causal conclusions should therefore be drawn cautiously. Furthermore, over-interpretation of effect size estimates should generally be avoided. Instead, MR analyses should focus primarily on identifying the existence and direction of causal pathways.

Ideally, any findings would be supported by background knowledge on the underlying biology and, whenever possible, a triangulation of evidence. The increasing availability of GWAS results on very large imaging cohorts and advances in the understanding of pathways from genetic variants to higher-level, imaging-related phenotypes in future will allow for more detailed, robust and reliable analyses of the causal relationships between genetic, clinical and imaging variables on one side and measures of health outcomes on the other. Mendelian randomisation techniques and other causal inference methods have the potential to play a valuable role in identifying and interpreting these putative causal relationships.

**Ethics Statement**

Informed consent was obtained from all UK Biobank participants. Ethical procedures are controlled by a dedicated Ethics Advisory Committee (http://www.ukbiobank.ac.uk/ethics).

**Data Availability**

UK Biobank data are available through an application process. GWAS data are openly available from the IEU Open GWAS Project at https://gwas.mrcieu.ac.uk/.

**Code Availability**

Supporting code for the example applications is available at https://git.fmrib.ox.ac.uk/ndcn1032/mrneuroimg.

## Acknowledgements

We thank Stephen Burgess from the University of Cambridge for valuable comments. This research has been conducted in part using the UK Biobank resource under Application Number 8107; we are grateful to all UK Biobank participants. This work was primarily supported by a Wellcome Trust Collaborative Award (215573/Z/19/Z). The Wellcome Centre for Integrative Neuroimaging (WIN FMRIB) is supported by core funding from the Wellcome Trust (203139/Z/16/Z).

## Supplementary Figures

**Figure 12:**
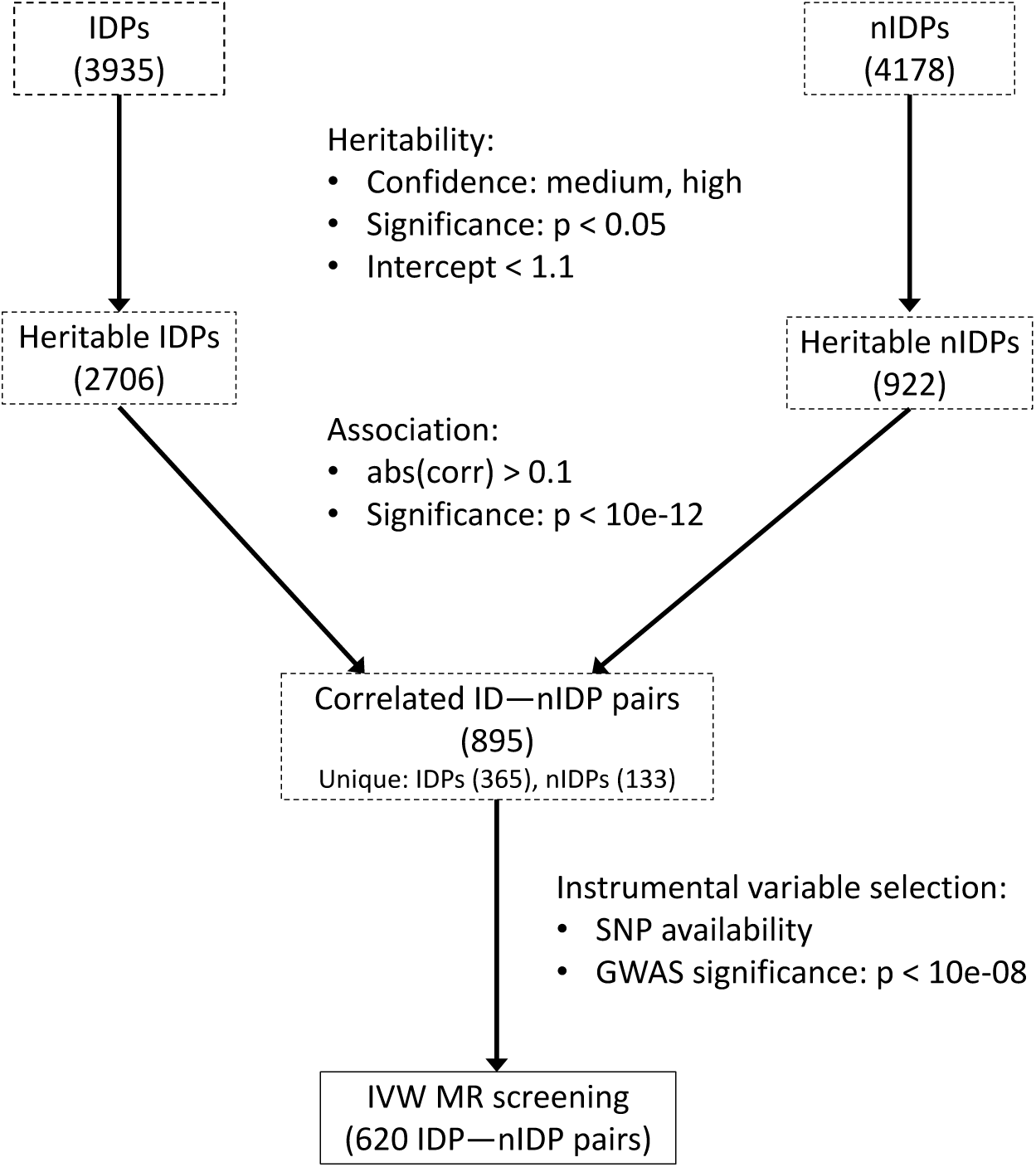
Flowchart showing the pre-selection and screening procedures of potential causal variable pairs as described in Subsections 3.1.1 and 3.1.2.

**Figure 13:**
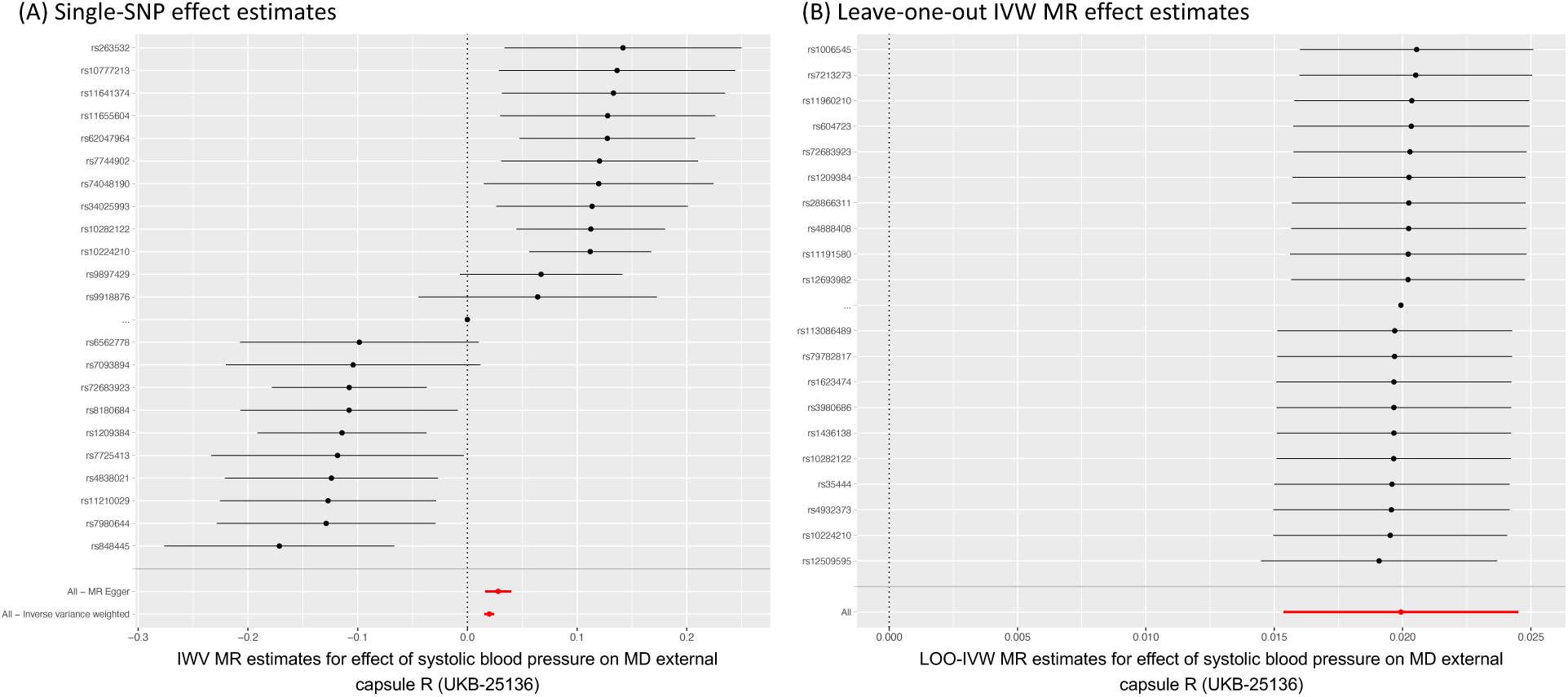
MR sensitivity analysis of the causal effect of systolic blood pressure on mean diffusivity of the external capsule WM tract R (UKB-ID 25316). (A) Single-SNP ratio estimates. (B) Leave-one-out MR results. Only results for 20 SNPs with the most extreme effect estimates are shown. Errorbars indicate 95% confidence intervals.

**Figure 14:**
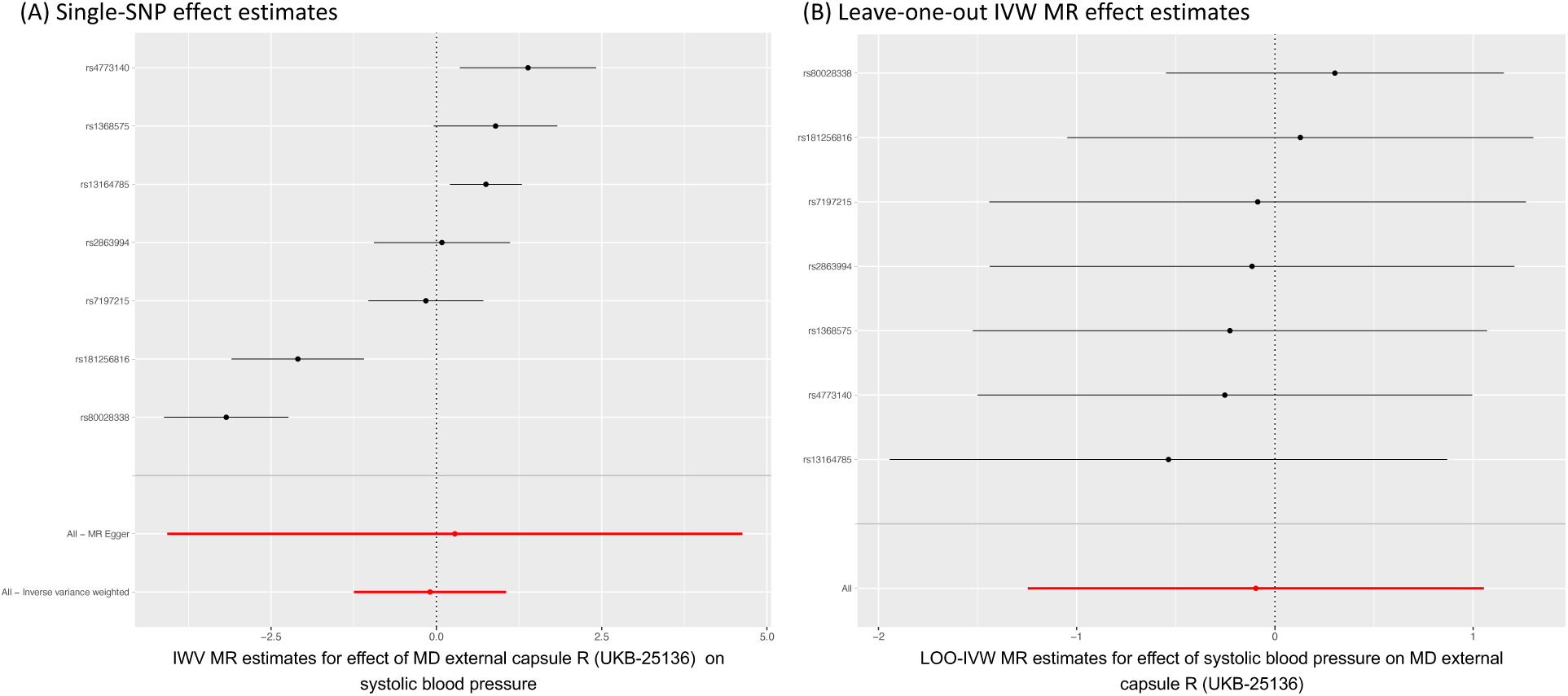
MR sensitivity analysis of the causal effect of mean diffusivity of the external capsule WM tract R (UKB-ID 25316) on systolic blood pressure. (A) Single-SNP ratio estimates. (B) Leave-one-out MR results. Errorbars indicate 95% confidence intervals.

**Figure 15:**
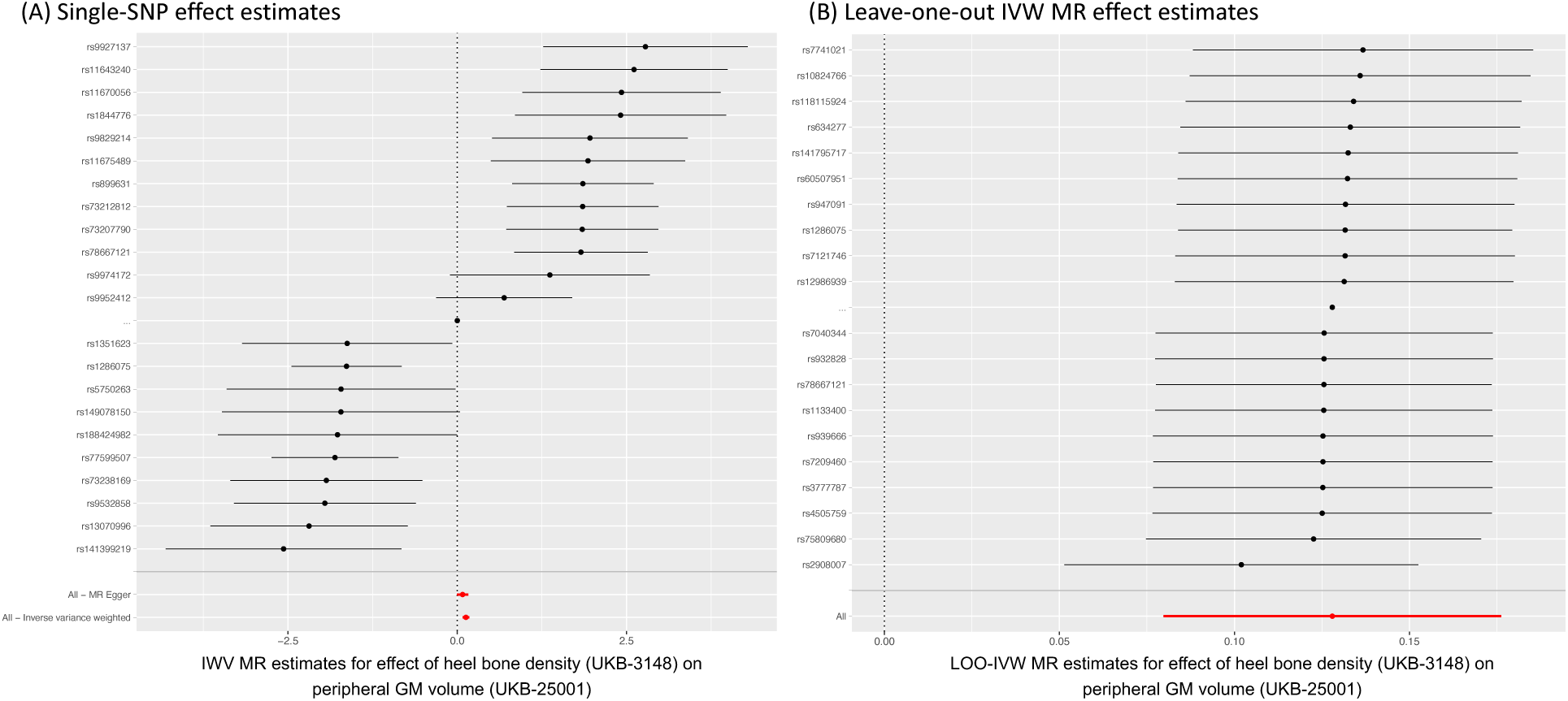
MR sensitivity analysis of the causal effect of heel bone density (UKB-ID 3148) on peripheral grey matter volume (UKB-ID 25001). (A) Single-SNP ratio estimates. (B) Leave-one-out MR results. Only results for 20 SNPs with the most extreme effect estimates are shown. Errorbars indicate 95% confidence intervals.

**Figure 16:**
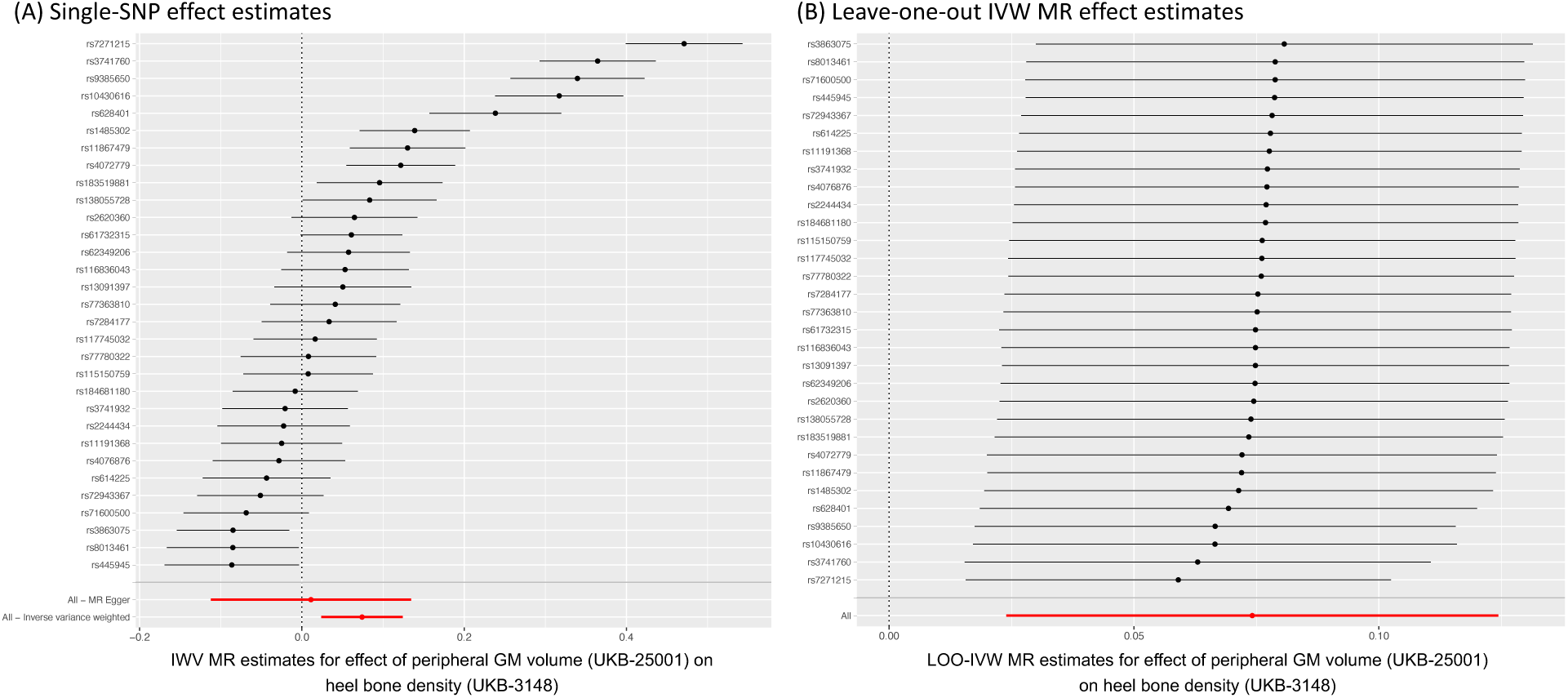
MR sensitivity analysis of the causal effect of mean diffusivity of peripheral grey matter volume (UKB-ID 25001) on heel bone density (UKB-ID 3148). (A) Single-SNP ratio estimates. (B) Leave-one-out MR results. Errorbars indicate 95% confidence intervals.

**Figure 17:**
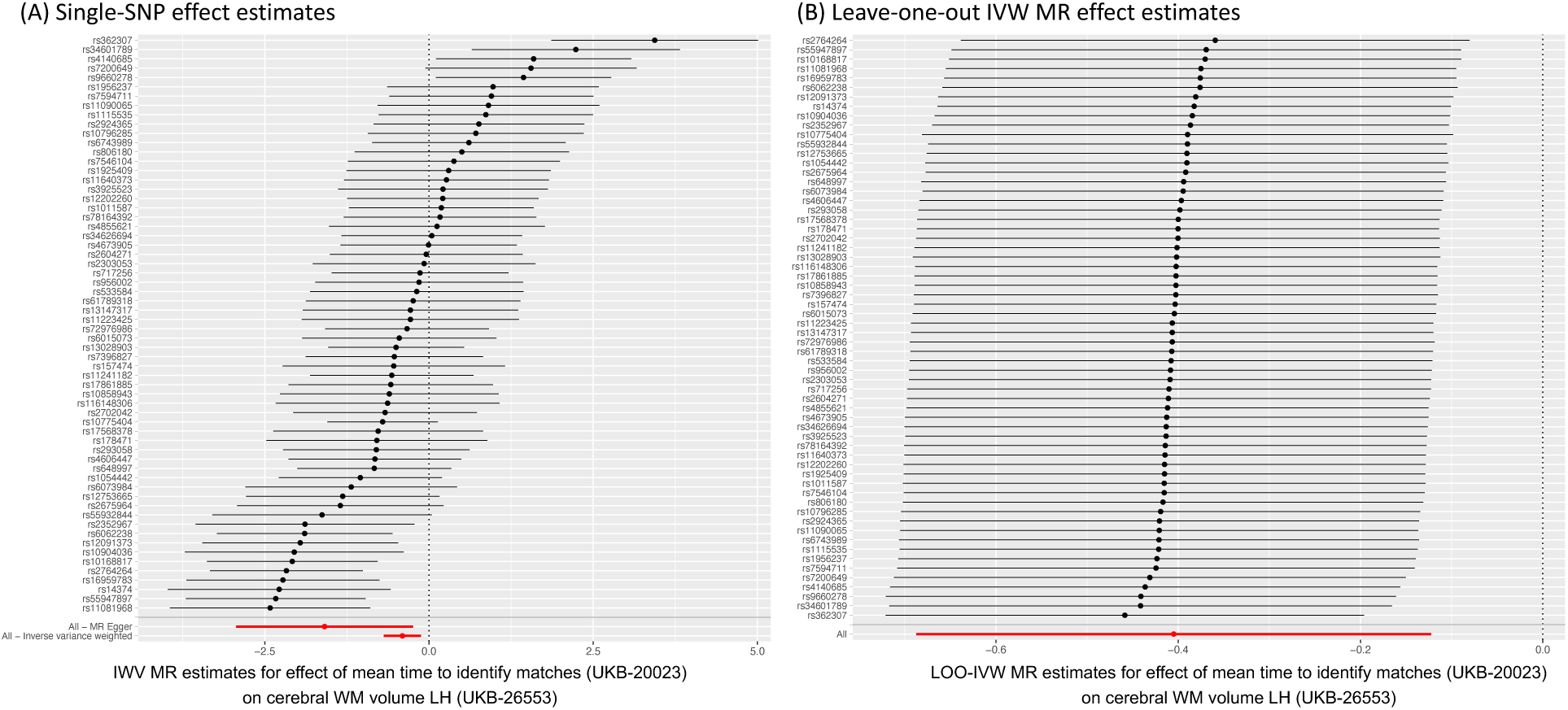
MR sensitivity analysis of the potentially causal effect of correctly identifying matches (UKB-ID 20023) on cerebral white matter volume (UKB-ID 26553). (A) Single-SNP ratio estimates. (B) Leave-one-out MR results. Errorbars indicate 95% confidence intervals.

**Figure 18:**
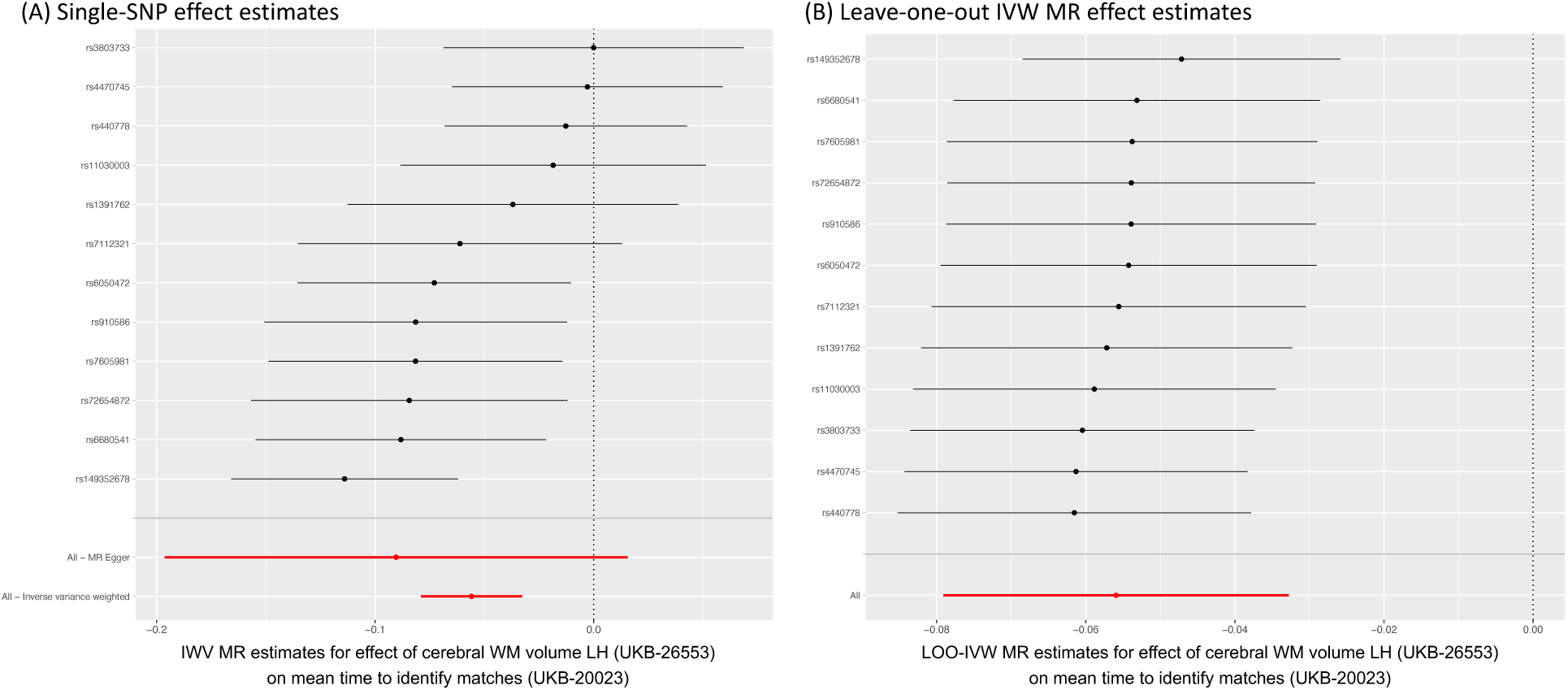
MR sensitivity analysis of the potentially causal effect of cerebral white matter volume (UKB-ID 26553) on correctly identifying matches (UKB-ID 20023). (A) Single-SNP ratio estimates. (B) Leave-one-out MR results. Errorbars indicate 95% confidence intervals.

## Footnotes

1 A confounder is a common cause of two or more variables, thereby introducing spurious correlations between these variables.

2 In economics and epidemiology, scenarios involving observational data and exposures outside the control of the investigator are also known as “natural experiments”.

3 Although the vast majority of MR studies uses SNPs as instrumental variables, other genetic variants such as indels and genetic variants associated with different gene expression or protein levels (eQTLs, pQTLs) can be used as instruments. For simplicity, we only refer to SNPs in this work.

4 For each SNP *j*, 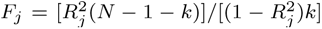, with GWAS sample size *N* , number of SNPs *k* and proportion of exposure variance explained by SNPs *R*^2^. In multivariable MR, the conditional F-statistic should be used instead.

5 This is analogous to the two-step least-squares (2SLS) approach for individual-level data, where in the first step the exposure is regressed on the SNPs and in the second step the outcome is regressed on the fitted values of the exposure.

6 In detail, this follows from the following linear relations (*X* denoting the exposure, *Y* the outcome and *Z* the SNP): *X* = *Zγ_E_* + *ɛ* and *Y* = *Zγ_O_* + *ɛ*, leading to *Y* = *Xβ* + *ɛ* = *Zγ_E_β* + *ɛ*, and finally *γ_O_* = *γ_E_β*. Rearranging and substituting sample estimates gives 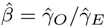.

7 Higgins’ *I*^2^ is defined in relation to Cochran’s *Q* as *I*^2^ = *Q* (*L* 1)*/Q*, where *L* denotes the number of SNPs. It can be used to assess regression dilution and expected bias (towards null) of the MR-Egger estimate.

8 The Neale lab results offer a confidence metric for each heritability result of None, Low, Medium or High, based on sample size, standard error, potential sex bias and other possible issues.

9 We note that all results should be interpreted with the caveat in mind that there exists an asymmetry in power due to sample size, meaning that MR analyses with IDPs as outcome (but not exposure) generally have higher power than MR analyses that involve IDPs as exposure.

10 Note that, although not present here, bi-modality in the MR-Mix output can be an indication that the wrong exposure– outcome direction has been specified. In the present example, a second peak away from zero would point to the existence of a non-zero effect in the forward direction.

## Notes

### Competing Interest Statement

The authors have declared no competing interest.

